# Engineering T cells to enhance 3D migration through structurally and mechanically complex tumor microenvironments

**DOI:** 10.1101/2020.04.21.051615

**Authors:** Erdem D. Tabdanov, Nelson J. Rodríguez-Merced, Alexander X. Cartagena-Rivera, Vikram V. Puram, Mackenzie K. Callaway, Ethan A. Ensminger, Emily J. Pomeroy, Kenta Yamamoto, Walker S. Lahr, Beau R. Webber, Branden S. Moriarity, Alexander S. Zhovmer, Paolo P. Provenzano

## Abstract

Defining the principles of T cell migration in structurally and mechanically complex tumor microenvironments is critical to understanding sanctuaries from antitumor immunity and optimizing T cell-related therapeutic strategies. To enhance T cell migration through complex microenvironments, we engineered nanotextured platforms that allowed us to define how the balance between T cell phenotypes influences migration in response to tumor-mimetic structural and mechanical cues and characterize a mechanical optimum for migration that can be perturbed by manipulating an axis between microtubule stability and force generation. In 3D environments and live tumors, we demonstrate that microtubules instability, leading to increased Rho pathway-dependent cell contractility, promotes migration while clinically used microtubule-targeting chemotherapies profoundly decrease effective migration. Indeed, we show that rational manipulation of the microtubule-contractility axis, either pharmacologically or through genome engineering, results in engineered T cells that more effectively move through and interrogate 3D matrix and tumor volumes. This suggests that engineering cells to better navigate through 3D microenvironments could be part of an effective strategy to enhance efficacy of immune therapeutics.

## INTRODUCTION

While chemical signals play a strong role in drawing T cells into solid tumors, the physical features of the stroma (e.g. architecture and mechanics) also strongly influence T cell infiltration, and their ability to effectively distribute throughout, and sample, the entire tumor mass. Indeed, the complex stromal reaction in solid tumors can limit access and effective distribution of T cells creating antitumor immunity-free sanctuaries (Elahi-Gedwillo et al., 2019; Hartmann et al., 2014; Kuczek et al., 2019; Salmon et al., 2012; Stromnes et al., 2017). Furthermore, many solid tumors are rich with aligned extracellular matrix (ECM) networks (Best et al., 2019; Provenzano et al., 2006; Ray et al., 2017a), which provide contact guidance for carcinoma cells (Provenzano et al., 2006, 2008; Tabdanov et al., 2018a, 2018b), and can also direct migration of infiltrated T cells in solid tumors (Hartmann et al., 2014; Salmon et al., 2012; Wolf et al., 2003). Yet, our understanding of how native and engineered T cells migrate through mechanically complex tumor microenvironments (TMEs) is quite incomplete. While considerable efforts have been made to better understand the biophysics of the immune synapse (Dustin et al., 2010; Tabdanov et al., 2015), much less is known about the regulation of T cell migration in physically complex microenvironments. However, elucidating migration behavior and employing rational engineering design approaches, with genome and cell engineering, to alter native cytotoxic T cells and/or further enhance engineered T cells so that they can most effectively migrate through and sample the entire tumor volume can enhance therapeutic efficacy against solid tumors and indeed further improve cell-based cancer therapeutics in general.

T cell 3D motility inside healthy and tumor tissues is influenced by dynamically balanced superposition of phenotypes, such as an amoeboid phenotype with pseudopodia and a more mesenchymal-like phenotype with spreading lamellipodia, a phenomenon known as amoeboid-mesenchymal plasticity (Gaylo et al., 2016; Talkenberger et al., 2017), where phenotypic switching can be advantageous for navigating heterogeneous cell and ECM conditions. Thus, understanding the key principles of the amoeboid-mesenchymal plasticity balance in T cells during normal tissue scanning and migration through tumors is critical for understanding T cell motility and enhancing cancer immunotherapy. As such, platforms that capture key physical features of ECM microenvironments that induce relevant phenotypes and drive phenotypic switching, or co-existent phenotypes, are critical for understanding of T cell migration. While, T cell motility has been studied on 2D systems (Saitakis et al., 2017) and in 3D microchannels (Holle et al., 2019; Jacobelli et al., 2010), to date, existing platforms have not captured tumor relevant ECM architectures, such as aligned fiber networks (Provenzano et al., 2006; Ray et al., 2017a; Tabdanov et al., 2018b), and variable stiffness to distinctly, yet simultaneously, capture coexisting amoeboid and mesenchymal modes of migration. Indeed, although flat 2D systems can have great utility for studying T cell motility (Krummel et al., 2016; Tooley et al., 2009), they lack the capacity to facilitate steric T cell-microenvironment interactions from textured and aligned ECMs (Tabdanov et al., 2018b) that can be associated with more amoeboid modes of T cell migration, and are distinct from microchannels where locomotion takes place under uniform spatial confinement (Jacobelli et al., 2010). Thus, engineered platforms that incorporate disease relevant ECM architecture, with variable stiffness, that allow high-throughput quantitative analysis of motility for large cell numbers, are needed in order to dissect out the fundamental drivers of diverse T cell migration modes. Likewise, evaluation in more complex 3D environments (e.g. tissue engineered 3D matrices and live tumor microenvironments) are also needed to test the impact of T cell engineering in order to generate the optimal amoeboid-mesenchymal plasticity balance for movement throughout solid tumors.

Here, we use engineered platforms to elucidate fundamental modes of T cell migration governed by ECM architecture and mechanics and establish that stiffness-dependent T cell guidance, from disease-relevant aligned ECM, is greatly enhanced from amoeboid behavior that competes against a coexisting mechanically-controlled mesenchymal-like mode. By disrupting microtubule (MT) dynamics or increasing cell contractility we show enhanced migration in complex microenvironments and demonstrate that this microtubule-contractility axis can be confirmed and captured by using CRISPR technology to genome engineer T cells lacking GEF-H1. Knockout of GEF-H1 in the context of microtubule instability abrogates the increased migration response, while GEF-H1 without MT disruption significantly enhances migration. We test our findings in engineered 2D to 3D systems and in live tumor pancreatic ductal adenocarcinomas (PDA), a cancer that has a robust fibrotic and immunosuppressive stroma (Clark et al., 2007; Olive et al., 2009; Provenzano et al., 2012; Stromnes et al., 2014), and is frequently characterized by very limited and/or heterogeneous distributions of cytotoxic T cells (Elahi-Gedwillo et al., 2019; Feig et al., 2013; Jiang et al., 2016; Stromnes et al., 2014). Yet, PDA can become susceptible to immune checkpoint blockade following approaches to specifically disrupt the stroma to increase T cell infiltration and function (Feig et al., 2013; Jiang et al., 2016), consistent with our conclusion that specifically engineering T cells more effectively navigate through complex TMEs can be part of an effective strategy to maximize the impact of T cell-related immunotherapeutic approaches.

## RESULTS

### Mechanically regulated T cell interactions with cancer-relevant nanoarchitectures

We previously demonstrated that aligned stromal ECM in tumors, which has been reported to guide tumor-infiltrating T cells (Hartmann et al., 2014; Salmon et al., 2012), contains nanopatterned spacing and textures that can be effectively mimicked through textured nanopatterning technology (Ray et al., 2017a; Tabdanov et al., 2018b). This allows us to impart relevant directional migration cues that are a useful compromise between two-dimensional (2D) and three-dimensional (3D) environments and allow us to perform quantitative analysis on large cell numbers under defined conditions. Therefore, in order to decipher how tumor architectures and mechanics influence T cell motility, we designed “2.5D” nanotextured platforms with defined mechanical rigidities and oriented architectures. These platforms are sterically interactive as they concurrently allow for both T cell amoeboid invasiveness (i.e. sterically-interactive pseudopodia) into textures and more mesenchymal-like “on-ridge” lamellipodia spreading behavior (**Figure 1 and Supplemental Figure 1**), where the mesenchymal-like phenotype for T cells is defined as lamellipodium-driven protrusions during migration, particularly on flat environments. To generate soft to stiff nanopatterned platforms we engineered high definition nanotextured polyacrylamide gel (PAAG) surfaces with Hookean elasticity and shear moduli of G’=2.3, 8.6, 16, 50, and ≫1,000 kPa (Tabdanov et al., 2018a, 2018b), and functionalized surfaces with either ICAM1 or ECM, and utilized these platforms to dissect architectural and mechanical regulation of T phenotypic plasticity and cell migration.

**Figure 1.**
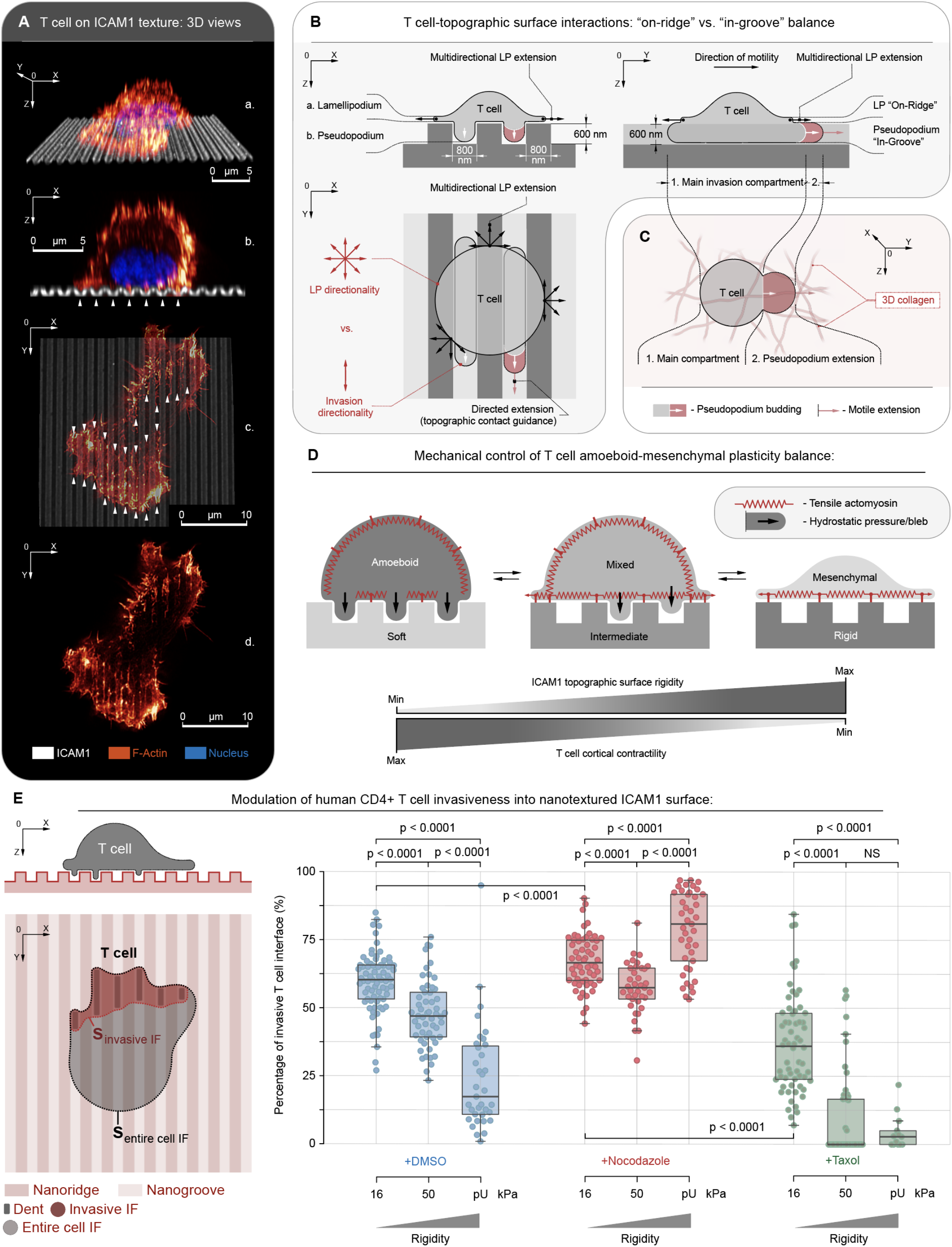
Mechanical and microtubule regulated T cell interactions with cancer-relevant nanoarchitectures (see also Supplementary Figure 1). **(A)** Super-resolution STED imaging used to generate 3D views of T cells on ICAM1-PAAG nanotopographies (G’=50 kPa in this example) in order to characterize T cell interactions with nanotopography. Views: a - stereometric, b - cross-section, c - interface from atop (*arrowheads* - “in-groove” invasions), and d - F-actin (phalloidin). **(B)** Projections schematics of nanotexture-T cell interactions, showing simultaneous “on-ridge” and “in-groove” dynamics. The nano-“ridge/groove” configuration supports both (a) unconfined, multidirectional “on-ridge” lamellipodium (LP), and (b) “in-groove” sterically interactive pseudopodium. “In-groove” T cell dentations are sterically confined and are guided along the nanogrooves, driving T cell contact guidance. **(C)** T cell motility in 3D extracellular matrix where extension of invasive pseudopodium, and rear retraction, propel T cells through the 3D environment. Although 3D pseudopodia may not be confined to singular motility direction, they remain analogous to directed “in-groove” dentations. **(D)** Mechanomodulated “in-groove” (i.e. more amoeboid-like phenotype with actin-rich pseudopodia) and “on-ridge” (i.e. more mesenchymal-like phenotype, which for T cells is defined as lamellipodia driven protrusions during migration, particular on flat environments) dynamics can be competitive and regulate the amoeboid-mesenchymal plasticity balance competition. Our model predicts “on-ridge” mesenchymal spreading enhancement on rigid nanotopography outbalances and antagonizes “in-groove” amoeboid invasiveness. Alternatively, soft nanotextures shift the balance towards amoeboid “in-groove” steric cell invasiveness. **(E)** Metrics (*left*) and measurement (*right*) of T cell “in-groove” relative invasiveness as a ratio between area of invasive regions and entire T cell area at the topography interface (i.e. 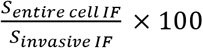). Invasiveness is a function of nanotopography rigidity G’ and microtubule stability. G’ range: 16 kPa to 50 kPa (PAAG) to ≫1,000 kPa (pU: polyurethane plastic). Remarkably, at ICAM1-PAAG nanotextures of G’<16 kPa hCD4+ T cells fail to attach to the nanotopographies, indicating G’≈16kPa may be the lower mechanosensing limit for hCD4+ T cells on ICAM1.

Examination of the interactions between activated primary human T cells and elastic nanotopography reveals a partial T cell invasiveness into the nanogrooves depth (**Figure 1A**), a level of the topographic interface termed the “in-groove” plane (**Figure 1B**). The remaining “noninvasive” part of the T cell is structured more like a flat lamellipodium spread atop of the nanotextured plane (i.e. more mesenchymal-like), termed the “on-ridge” level (**Figure 1B**). This is consistent with carcinoma cell behavior where competitive dynamics between nanogrooves-guided directed protrusions and less oriented “on-ridge” spreading influence cell orientation and migration (Tabdanov et al., 2018b), and consistent with electron microscopy findings of T cell lamellipodia spreading on top of nanostructured surfaces (Chen et al., 2014; Kwon et al., 2012). In our model, we draw parallels between invasive “in-groove” pseudopodia (**Figure 1A**) and T cell pseudopodial protrusions observed during migration through 3D collagen matrices (Wolf et al., 2003), and to the amoeboid locomotion reported within and along the uniaxially confining microchannels (Holle et al., 2019; Jacobelli et al., 2010) (**Figure 1B,C**). However, since the nanogroove platforms developed here facilitate partial confinement that allows for both highly directed “in-groove”-confined amoeboid invasions and less directed “on-ridge” 2D mesenchymal-like lamellipodium protrusions (**Figure 1A,B**), i.e. a superposition of amoeboid and mesenchymal functional phenotypes (**Figure 1B**), they allow us to define the mechanical regulators that drive the amoeboid-mesenchymal plasticity balance (**Figure 1D,E**) and define ways for us to tilt the balance in order to design T cells that can more effectively move throughout TMEs.

From a mechanobiology perspective, “in-groove” pseudopodia formation is primarily initiated by cytoplasm hydrostatic pressure (Ruprecht et al., 2015) generated by cortical actomyosin contractility. Cytoplasm pressure-driven off-cortex plasma membrane peeling (Charras et al., 2005, 2006; Tyson et al., 2014) results in budding of the invasive plasma membrane blebs, i.e. pseudopodial precursors (**Figure 1B**), which then rebuild an actomyosin cortex in the new pseudopodia extensions (Charras et al., 2005, 2006; Tyson et al., 2014). Therefore, actin-rich pseudopodia formation dynamics can be inherently antagonized by mechanical tension from the more mesenchymal behavior within the “on-ridge” cell-substrate adhesion interface (**Figure 1B,D**), similar to observations in carcinoma cells (Tabdanov et al., 2018b). This suggests that the mechanical properties of the substrate, and the related intracellular mechanics, play a role in governing directed motility of T cells. To explore this regulation, we first engineered nanolines (flat 2D lines) and nanotextures (“2.5D”) of different stiffnesses functionalized with ICAM1. Evaluation of activated human T cell dynamics demonstrates mechanoresponsive behavior with significantly decreased “in-groove” interactions (i.e. “in-groove” invasiveness) as nanotexture stiffness increases (**Figure 1E**). Indeed, increasing mechanical rigidity from G’=16 kPa through 50 kPa (polyacrylamide gel: PAAG) to G’≫1 GPa (polyurethane, i.e. pU) results in a significant and consistent drop of “in-groove” invasiveness (defined as the invasive area percentage of cell-substrate interface projection) from ~60% to ~45% to ~20%, respectively (**Figure 1E; Supplemental Figure 1A-C**). Thus, our data suggest that partial T cell “in-groove” amoeboid invasiveness is regulated by mechanotransduction resulting from differences in the mechanical rigidity of the microenvironment. This is consistent with findings that ICAM1 or VCAM1 substrate stiffness influences T cell behavior (Saitakis et al., 2017; Schaefer and Hordijk, 2015) and regulation of T cell actomyosin tension at cell-substrate adhesions by a LFA1 mechanosensing feedback pathway (Nordenfelt et al., 2016; Walling and Kim, 2018), and thus we propose that increasing rigidity of nanotextures (control, i.e. +DMSO) results in an increase in T cell traction at the cell-substrate interface that antagonizes “in-groove” invasive pseudopodia.

### Microtubules regulate the amoeboid-mesenchymal plasticity balance

We previously demonstrated that MT-rich protrusions provide a scaffold that sterically and mechanically enhances “in-groove” invasiveness into “2.5D” surfaces to promote cell alignment and directed migration of carcinoma cells, i.e. contact guidance (Tabdanov et al., 2018b). Furthermore, Nocodazole treatment has been shown to selectively increase T cell blebbing and enhanced pseudopodia-like shape shifting dynamics (Takesono et al., 2010), which we link here to the increased T cells invasive interactions with nanotexture contact guidance cues. Therefore, we hypothesized that MTs may play a role in regulating T cell interactions with nanoarchitectures and the degree of the resulting amoeboid and mesenchymal-like behaviors. To test a role for MTs in regulating phenotype and associated migration in human T cells, we pharmacologically destabilized MTs with Nocodazole or stabilized MTs with Taxol and evaluated resulting “in-groove” invasiveness. Perhaps surprisingly, in contrast to carcinoma cells, MT destabilization (+Nocodazole) profoundly increases T cell amoeboid “in-groove” invasiveness (area fraction>50%) across all nanotexture mechanical rigidities (**Figure 1E and Supplemental Figure 1; +Nocodazole**). In stark contrast, MT stabilization with Taxol induced a universal and substantial reduction of “in-groove” invasiveness for each substrate stiffness (area fraction<10-35%) without the loss of “on-ridge” T cell adhesion and spreading across any of the “2.5D” substrate rigidities (**Figure 1E and Supplemental Figure 1; +Taxol**). Hence, altering MT abundance or dynamics changes the fundamental interaction between T cells and textured surfaces that mimics *in vivo* textures and thus T cell phenotype. This suggests that MT state may influence T cell migration, which can inform strategies to enhance T cell migration in tumors, and also suggests that clinically utilized drugs such as Taxol could be detrimental to T cell migration.

### Microtubule dynamics regulate substrate texture- and stiffness-dependent directed migration

To define the impact of stiffness on directed T cell migration, we analyzed cell directionality relative to nanotopography (i.e. contact guidance) and migration speed on aligned substrates with stiffnesses ranging from 16 kPa to over 1 GPa (**Figure 2A-C**); a range were we observed superpositioned phenotype behavior with “in-groove” amoeboid-like and “on-ridge” mesenchymal-like dynamics in response to the “2.5D” nanoarchitectures (**Figures 1 and 2D**). By tracking cell movement over time, we identified distinct differences in T cell contact guidance between soft and rigid conditions. On lower modulus nanotexture conditions, T cell migration is strongly contact guidance-driven with highly oriented migration along the direction of the nanogrooves (**Figure 2B**). In contrast, cells on stiffer substrates display more random migration tracks relative to the directionality of the underlying nanogrooves (**Figure 2B**). This demonstrates that the shift toward a more amoeboid phenotype regulates migration in structurally complex environments in a substrate stiffness-dependent manner (**Figure 2D**). Indeed, quantification of T cell migration directionalities per every 10 second-step across distinct nanotexture rigidities (16 to 50 kPa to >1,000 kPa) demonstrates significant differences in T cell migration directionality, with collectively stronger guidance along the 0° angle of deviation from the nanogrooves orientation on soft (G’=16 kPa) nanotextures, but more random and less persistent directionalities on stiffer nanotextures **(Figure 2E**). Furthermore, as expected from our analysis of T cell “in-groove” vs. “on ridge” interactions with nanotexture (**Figure 1**), in control conditions (+DMSO), T cell migration displays a direct and proportional correlation to the degree of T cell “in-groove” invasiveness (i.e. the amoeboid-mesenchymal phenotype balance), with significantly slower migration as substrate stiffness increases (i.e. shifted toward a larger contribution from the “on-groove” mesenchymal phase; **Figure 2A-C, +DMSO**). This again shows that softer environments more robustly facilitate fast directed migration of T cells resulting from a shift toward a larger contribution from the “in-groove” amoeboid phase. This also suggests that nanotextures are well-suited for steric interactions-driven amoeboid-like motility via “in-groove” T cell pseudopodia dynamics that facilitate a high-speed mode of migration. Furthermore, importantly, the principles of the mechanically-regulated amoeboid-mesenchymal balance that we report here and its effects on T cell directed migration are not limited to interactions with ICAM1, as we observe similar trends for T cell migration along fibronectin (FN) nanotextures of the same mechanical rigidity range (**Supplemental Figure 2A,B**). Again directionality and speed are maximal on softer FN nanotextures (**Supplemental Figure 2A,B**), suggesting that the architecture and mechanics of the environment are dominant factors that can drive T cell motility across a range of complex ECM environments. These cumulative data also suggest that the increased tissue stiffness associated with TMEs (vs. normal tissues) may in fact reduce T cell migration and be part of the physical barrier to effective T cell sampling in tumors.

**Figure 2.**
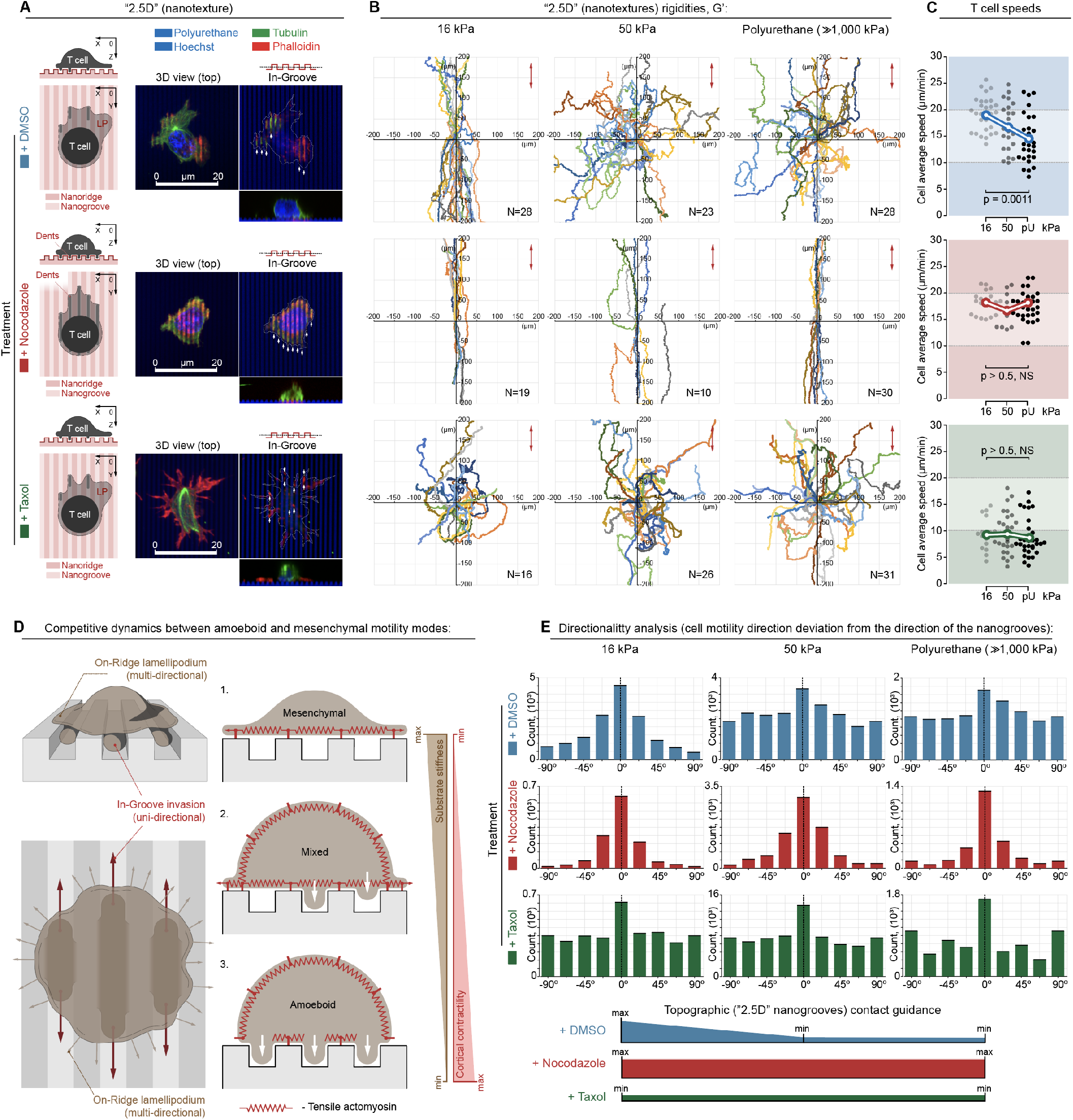
Microtubule dynamics regulate texture- and stiffness-dependent directed migration (see also Supplementary Figure 2). **(A)** *Left to right*: Schematic, 3D micrograph, and “in-groove” cross-section and side views of human T cells on ICAM1 nanotextures (grooves/ridges widths = 800 nm). *Top to bottom*: control (+DMSO), destabilized MTs (+Nocodazole), and stabilized MTs (+Taxol) showing increased “in-groove” actin-rich pseudopodia in cells with destabilized MTs, in contrast to more mesenchymal-like behavior with Taxol. **(B)** T cells migration tracks on compliant (G’=16 kPa), intermediate (G’=50 kPa), or rigid (G’≫1,000 kPa) ICAM1 nanotextures, where compliant nanotopographies enhance contact guidance. (*Top to bottom*) T cells migration under control (+DMSO), Nocodazole, or Taxol treatment conditions, where MT disassembly results in the enhanced directed migration across all rigidities. **(C)** T cell averaged per cell speeds on ICAM1 nanotextures of various rigidities (16, 50 and ≫1,000 kPa) for control (+DMSO, *top*), Nocodazole, (*middle*), Taxol (*bottom*) conditions. **(D)** *Left* - Schematics of T cell-ICAM1 nanotexture interactions. *Right* - Mechanosensitive “in-groove” steric invasion and “on-ridge” mesenchymal-like (lamellipodial-based) interactions as a function of the mechanical rigidity of the microenvironment and cell contractility. **(E)** Quantification of directionality of human T cells migration during contact guidance as a function of substrate mechanical rigidity and the state of microtubules. Corresponding T cells migration tracks are shown in the same matrix order in panel B. Measurements represent frequency distributions of cell-to-nanogroove angles every 10 s step.

Remarkably, we note that our observed mechanoresponsive T cell behavior during interactions with disease-relevant nanotextures is not recapitulated on flat environments (**Figure 3**). Using anisotropic ICAM1 nanolines, featuring the same lines/gaps dimensionality as “2.5D” nanoridges, i.e. 800 nm-wide lines alternating with 800 nm-wide gaps, we observe T cell behavior that is distinct from behaviors on nanotextured surface and dependent on phenotype (**Figures 2 and 3**). On flat surface with moduli ranging from 8.6 to 50 kPa (we note that below 8.6 kPa human T cells displayed no significant adhesion whereas beyond 50 kPa we do not observe any principal differences in behavior), T cell migration is largely random (i.e. meandering) with no significant migration alignment to the anisotropically oriented flat lines (**Figures 3A, B**). Interestingly, in this context, we observed a range of morphologies and identified coexisting amoeboid (more spheroid) and mesenchymal-like (more flattened/spread) phenotypes. Thus, the flat 2D nanolines system allows us to isolate phenotype behaviors that co-exists along a phenotype spectrum in response to nanotextures, albeit with the latter more accurately mimicking natureal ECM architectures. As the amoeboid-mesenchymal transition was sporadically frequent, we could not build separate continuous migration tracks over time for each of the cell phenotypic states. However, we quantified migration speed during each phenotypic state, where measuring distinctly amoeboid vs. mesenchymal migration speeds for individual T cells indicates phenotype-specific behavior (**Figure 3C**). In the more mesenchymal-like state, migration increases with substrate stiffness (speed goes from 6 to 7.5 to 10 μm/min for G’ of 8.6 to 16 to 50 kPa, respectively), consistent with behavior reported for T cells on flat non-patterned ICAM1 surfaces, where the distinction between mesenchymal vs. amoeboid T cell phenotypes was not made (Saitakis et al., 2017). In contrast, we defined the opposite trend in speed as a function of substrate stiffness for T cells in the amoeboid phenotype, where average speed decreases from ~15 to ~10 μm/min as nanoline rigidity increases (**Figure 3C**), consistent with observations on nanotextures where amoeboid behavior is a more favorable mode of migration (**Figure 2**). Thus, since both T cells’ “in-groove” invasiveness and T cell contact guidance along nanogrooves display related behavior to topography mechanical rigidity (**Figures 1 and 2**), we suggest a mechanistic model for efficient T cell directed migration. In this model, “in-groove” invasive pseudopodial protrusions sterically guide and drive T cell migration along the nanogrooves (**Figure 2D**), similarly to the established amoeboid-like leader bleb-guided cell migration scenario (Logue et al., 2015). Conversely, co-existing “on-ridge” mesenchymal lamellipodium antagonizes efficient contact guidance by promoting multidirectional T cell spreading. Indeed, since the balanced competition between “in-groove” guidance and multidirectional “on-ridge” migration is dynamically controlled by the mechanical rigidity of aligned nanotextures, T cell amoeboid directed migration from microenvironmental architecture becomes a mechanobiological function of stiffness of the environment. Thus, this provides the basic principles governing amoeboid-mesenchymal plasticity control and design rules to produce more efficient migration in mechanically complex environments.

**Figure 3.**
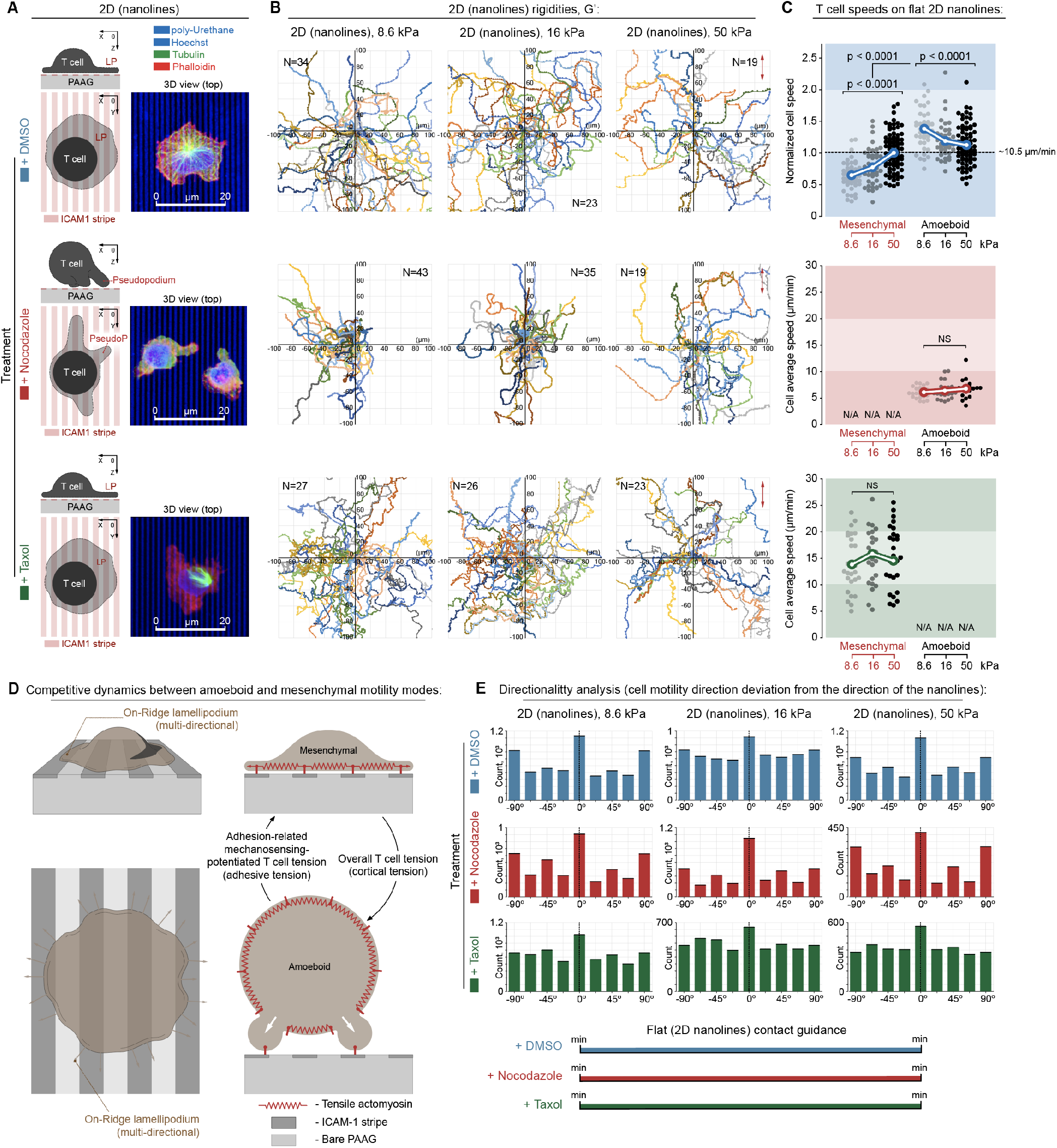
The microtubule-dependent amoeboid-mesenchymal phenotype balance influences T cell migration and mechanoresponsive behavior. **(A)** *Left to right*: Schematic, 3D micrograph, and cross-section and side views of human T cells on flat ICAM1 nanolines (line/gap widths = 800 nm). *Top to bottom*: control (+DMSO), destabilized MTs (+Nocodazole), and stabilized MTs (+Taxol) showing distinct MT stability-dependent phenotypes. **(B)** T cells migration tracks on compliant (G’=16 kPa), intermediate (G’=50 kPa), or rigid (G’≫1,000 kPa) flat ICAM1 nanolines. (Top to bottom) T cells migration under control (+DMSO), Nocodazole, or Taxol treatment conditions. **(C)** T cell averaged per cell speeds on ICAM1 nanolines of various rigidities (16, 50 and ≫1,000 kPa) under control (+DMSO, top), Nocodazole, (middle), Taxol (bottom) conditions showing that T cells in the flat control conditions (+DMSO) can display either amoeboid and mesenchymal phenotypes that result in different migration speeds and different mechanoresponsive behaviors, while MT instability shifts T cells to amoeboid behavior and MT stability shifts to the mesenchymal-like behavior. **(E)** Quantification of directionality of human T cells migration on flat nanolines as a function of substrate mechanical rigidity and the state of microtubules. Corresponding T cells migration tracks are shown in the same matrix order as panel B. Measurements represent frequency distributions of cell-to-nanoline angles every 10 s step.

Since destabilization of microtubules with Nocodazole induces a strong T cell phenotype shift towards amoeboid “in-groove” invasiveness across all “2.5D” substrates rigidities, (**Figure 1E and Supplemental Figure 1**), and since we established a link between amoeboid “in-groove” interactions and highly directed and rapid T cell migration (**Figures 2 and Supplemental Figure 2**), we hypothesized that disruption of MTs would enhance T cell migration across a range of mechanical environments. Indeed, MT disruption results in highly directed migration from contact guidance across all substrate stiffnesses on both ICAM and FN (**Figure 2A,B and Supplemental Figure 2A,B**). Likewise, under MT destabilization cell speed remains constant regardless of substrate stiffness (**Figure 2A,B and Supplemental Figure 2A,B**), indicating a loss of mechano-responsive behavior, and that following the shift toward the more amoeboid phenotype T cells are now able to migrate as efficiently in stiff environments as in soft environments. Interestingly, MT disruption also impacts the response to flat nanolines by severely limiting development of mesenchymal behavior while still resulting in no change in speed in response to increasing stiffness, albeit with a lower overall speed than the amoeboid state under control conditions (**Figure 3A-C**), which is consistent with our hypothesis that mechano-responsive behavior in response to nanotexture is regulated by competition from mesenchymal-like phase that competes with more efficient directed migration from guiding pseudopods. In stark contrast to enhanced migration along nanogrooves following robust MT destabilization, MT stabilization with Taxol induced the opposite trends. Taxol-treated T cells showed a substantial reduction of “in-groove” invasiveness (area fraction<10-35%) and significant decreases in directionality and speed (**Figure 2A-C**) indicating a shift from amoeboid phenotype toward the mesenchymal “on-ridge” T cell spreading, which is supported by findings on flat 2D substrates, where Taxol treated cell shift toward the more mesenchymal phenotype (**Figure 3B-E**). Thus, our results on “2.5D” environments indicate that commonly utilized microtubule stabilizing taxane chemotherapeutics may significantly reduce T cell migration, suggesting that analyzing migration of MT-stabilized T cells in tumor environments is needed, as caution may be warranted when timing chemotherapy and immunotherapy combination approaches that require T cell infiltration into tumors. Moreover, our data indicate that more destabilized MTs promote T cell migration in architecturally and mechanically relevant “2.5D” environments and offer insight into design criteria for approaches to enhance T cell migration in more complex 3D environments using pharmacological agents or via genome engineering. Thus, our data suggest that this approach could be applied to further impove therapeutic T cells (e.g. enhance cells that are already engineered with high affinity tumor-reactive TCRs specific to tumor antigens).

### Controlling T cell migration in 3D environments through the Microtubule-Contractility axis

We next examined how our findings that disruption of microtubules shifted activated human T cells toward enhanced migration due to more amoeboid-like behavior (**Figure 1, Figure 2 and Supplemental Figure 2**) would translate to more complex 3D conditions (**Figure 4A**). In 3D fibrous ECM networks with more random ECM organization, cells move as a series of short contact guidance steps along fibers, or groups of fibers, with the need to re-orient in order to transition to another fiber configuration for guidance cues, during tortuous paths in the network, where they must also go through pores or across fibers in dense ECM regions. In this context, we hypothesized that MT destabilization is influencing Rho signaling behavior via GEF-H1 (**Figure 4B**). We attribute our observed phenotype under MT destabilized condition to release of MT-sequestered GEF-H1, which activates the RhoA pathway that directly enhances downstream actomyosin contractility (Birkenfeld et al., 2007; Heck et al., 2012; Krendel et al., 2002; Ren et al., 1998). Therefore we tested our hypothesis by pharmacologic MT destabilization (+Nocodazole), control of GEF-H1 cytoplasmic levels, and by perturbing RhoA activity and downstream actomyosin contractility in order to manipulate the MT-contractility axis to control the amoeboid-mesenchymal balance (**Figures 4B,C**), where these factors regulates pseudopodia protrusion dynamics and later cell tractions as the cell moves through the 3D network (**Figure 4A,C**).

**Figure 4.**
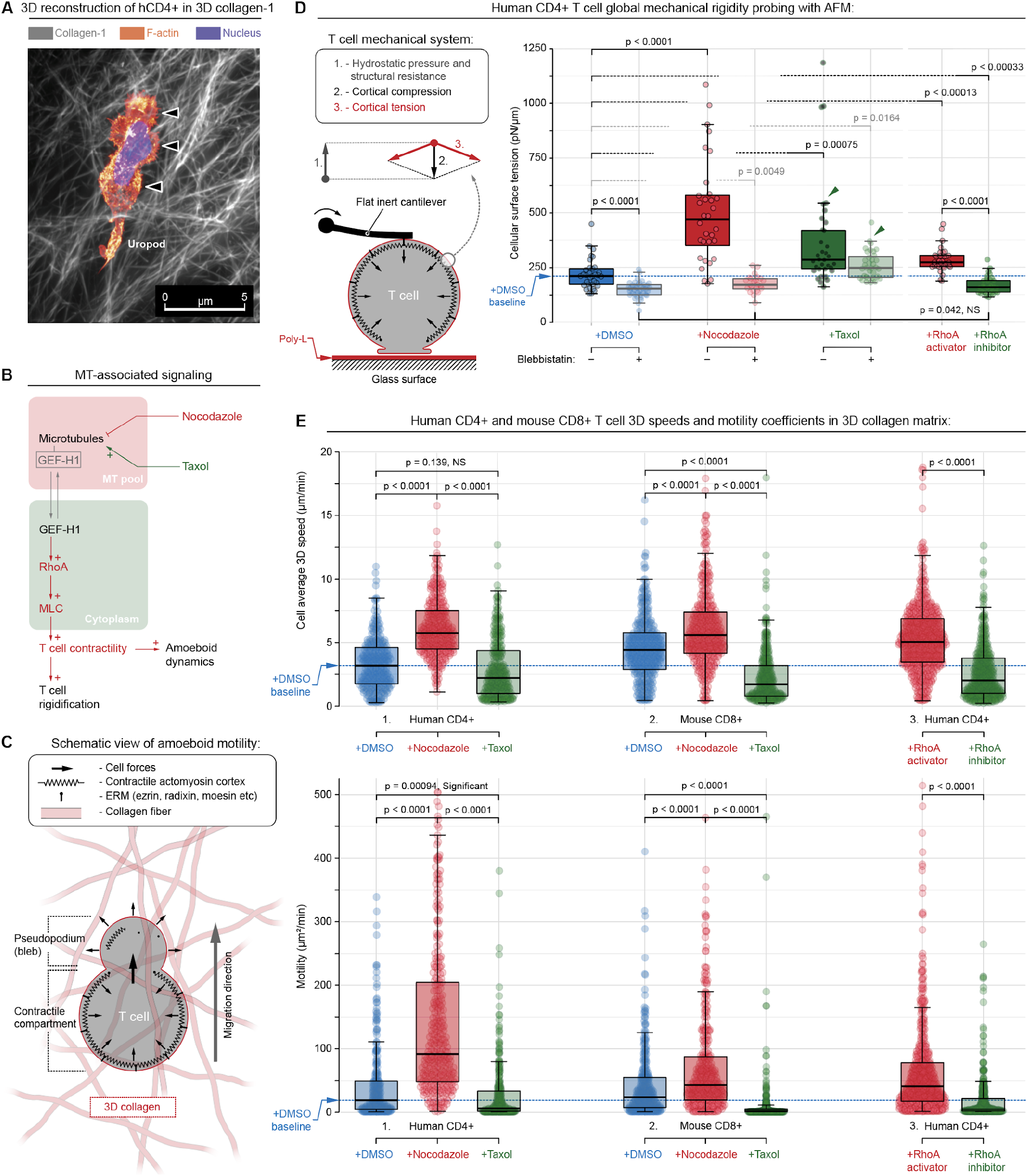
The microtubule-contractility axis regulates T cell mechanics and migration in 3D environments. **(A)** STED micrograph of a human T cell migrating through 3D collagen matrix. *Arrowheads* - distinct bleb-to-pseudopodia amoeboid protrusions. **(B)** Model for how MT-associated signaling integrates amoeboid motility through MT stability and contractility via GEF-H1 “cytoplasm⇄MT” reversible transitioning. **(C)** T cell amoeboid motility in 3D environments is linked to actomyosin contractility where cortex tension-induced hydrostatic pressure forces plasma membrane off-cortex peeling (blebbing) that results in pseudopodia formation. Continuous pseudopodial formation facilitates steric interactions between T cells and the 3D microenvironment, i.e. persistent amoeboid motility. **(D)** *Left* - Mechanical testing of T cells with atomic force microscopy (AFM) to measure to global compression through a flat cantilever to measure the overall cortical tension that is counterbalanced with cytoplasm hydrostatic pressure **(see also Supplemental Figure 3)** and additional intracellular cytoskeletal structural resistance (microtubules, nucleus etc). *Right* - AFM measurement of global T cell surface tension, a measure of cortical contractility. The control group and each of the two MT-targeting treatments (i.e., +DMSO, +Nocodazole and +Taxol, *solid colors*) are paired with blebbistatin co-treatment (*pale semi-transparent colors*) to verify the key role of actomyosin contractility as a regulator of changes in cell rigidity during MT-targeting. Alternatively, direct RhoA activation or inhibition is compared to MT-perturbation results. Both MT disassembly (+Nocodazole) and direct RhoA activation (+RhoA activator) induce mechanical rigidification of human CD4+ T cells via increased actomyosin tension, as demonstrated by AFM findings after blebbistatin co-treatment. Note, MT destabilization or direct RhoA activation induce a rise of hydrostatic pressure (**see Supplemental Figure 3**). Note, Taxol-induced MTs stabilization increases passive (i.e. actomyosin-independent) T cell rigidity via a direct mechanical contribution from stabilized MT scaffolds (+Taxol vs. +Taxol+Blebbistatin treatments; arrowheads). **(E)** 3D migration speed 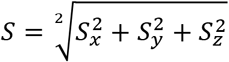, *top*) and overall 3D cell motility (*μ* = *μ_x_* + *μ_y_* + *μ_z_*, *bottom*) of T cells in 3D collagen-FN matrix: 1 - human T cells (hCD4+) in the presence of vehicle (+DMSO), Nocodazole (+Nocodazole) or Taxol (+Taxol); 2 - murine cytotoxic T cells (mCD8+) in the presence of vehicle (+DMSO), Nocodazole (+Nocodazole) or Taxol (+Taxol), and 3 - human T cells in the presence of RhoA activator (+RhoA activator) or inhibitor (+RhoA inhibitor).

To first explore the implications of manipulating MT stability and the overall microtubule-contractility axis, we employed atomic force microscopy (AFM) (Cartagena-Rivera et al., 2016) on primary human T cells under various pharmacologic conditions to alter MT stability or Rho contractility (**Figure 4D**). Here the corresponding actomyosin cortical tension contractile forces are balanced by opposite forces generated by the intracellular “cytosolic” hydrostatic pressure (Cartagena-Rivera et al., 2016; Ramanathan et al., 2015; Stewart et al., 2011). Importantly, the cellular cortical tension and the accompanying intracellular hydrostatic pressure concomitantly regulate the generation of cellular blebs and ameboid migration in complex 3D microenvironments (Charras et al., 2005; Tozluoğlu et al., 2013). Consistent with our model, results indicate that along with the universal shift towards a more amoeboid-like phenotype (**Figure 2 and Supplemental Figure 2**), MT destabilization significantly increases the mechanical rigidity as a function of actomyosin contractility, which is confirmed by treating cells with Blebbistatin to robustly disrupt myosin-regulated contractility (**Figure 4D, +DMSO vs. +Nocodazole**). In fact, we note the highly consistent T cell softening without myosin-based contractile forces to the nearly identical values observed during treatment with +Blebbistatin versus +raNocodazole+Blebbistatin (**Figure 4D**). Furthermore, consistent with our cortical tension observations, MT destabilization significantly increases the intracellular hydrostatic pressure, while Blebbistatin treatment reduced hydrostatic pressure (**Supplemental Figure 3**). Interestingly, Taxol also induces T cell rigidification and increased hydrostatic pressure, however, this stiffening was more independent of actomyosin contractility, as addition of Blebbistatin did not soften cells or reduce pressure as profoundly as in control or Nocodazole treatment conditions (**Figure 4D, Supplemental Figure 3**). We attribute this significant effect to rigidification via Taxol-stabilized microtubules (Cheng and Donhauser, 2013; Elie-Caille et al., 2007; Sun et al., 2009) and may also be related to findings that the microtubule cytoskeleton can physically connect and interact with the actomyosin cortex providing an additional structural scaffold (Dogterom and Koenderink, 2019; Rodriguez et al., 2003), which is also strongly in line with the recently identified contribution of scaffolding microtubules in active cell mechanics during directed migration (Tabdanov et al., 2018b). Lastly, direct RhoA activation was performed with G-switch™ treatments, which activates Rho by converting glutamine-63 to glutamate in the Switch II region in order to constitutively activate Rho, while inhibition was performed with cell-permeable C3 exoenzyme. Again, consistent with our hypothesis, activation of Rho increases T cell rigidity and hydrostatic pressure, while Rho inhibition softens cells and reduces hydrostatic pressure to levels observed following treatment with Blebbistatin (**Figure 4D, Supplemental Figure 3**). Collectively, this suggests that amoeboid-mesenchymal plasticity balance is indeed regulated by a microtubule-contractility axis, and that together with cortical tension, modulation of the intracellular hydrostatic pressure allows T cells to alter their shape, control their locomotion, and govern their mechanics while infiltrating through complex intratumoral microenvironments.

Equipped with this new understanding of T cell mechanics and their implications on T cell motility, we sought to evaluate and enhance T cells migration in more complicated 3D environments. Using 3D collagen-FN matrices we first quantified T cell speed, as well as overall motility using the persistent random walk model, for both primary human CD4+ (hCD4+) T cells and mouse CD8+ (mCD8+) cytotoxic T cells, where cytotoxic T cells were isolated from genetically engineered *KPC* mice bearing pancreatic tumors, which are highly faithful to the human disease (Elahi-Gedwillo et al., 2019; Hingorani et al., 2005; Provenzano et al., 2012). In both human and mouse T cells, MT destabilization with Nocodazole significantly increases speed by ~50-100% and overall motility ~2-4 fold (**Figure 4E**), again suggesting that MT destabilizing agents have the potential to enhance T cell migration in tumor-like architectures. Likewise, similar strategies may be employed to enhance Rho activation as the increase in migration observed in human T cells, following Rho activation, is consistent with increases following MT destabilization, while inhibition of Rho again decreases speed and overall motility (**Figure 4E**). Thus, manipulating the microtubule-contractility axis has the potential to significantly enhance T cell migration through ECM architectures found in tumors and therefore may provide a strategy that could be developed to enhance migration of therapeutic T cells (e.g. CAR-T cells or other engineered T cells) or native T cells during immunotherapy (e.g. Immune checkpoint blockade). To further test this hypothesis we isolated cytotoxic mCD8+ T cells from *KPC* mice harboring autochthonous metastatic pancreas cancer and imaged their migration over hours in paired live tumor slices (**Figure 5A**), with or without MT destabilization. Consistent with data on “2.5D” and in engineered 3D environments, microtubule disruption, specifically in T cells, significantly increased migration of cytotoxic T cells through native tumor environments (**Figure 5B**), suggesting that perturbing the MT-contractility axis is indeed a viable strategy that warrants development to enhance migration of native or therapeutic T cells though structurally complex tumor microenvironments in future studies. In stark contrast, T cell exposure to the MT stabilizing agent Taxol results in a significant decrease in both speed and overall motility both in 3D matrices *in vitro* (**Figure 4E**) and in live tumor (**Figure 5B**). This suggests that while Taxane agents can have a beneficial therapeutic impact on carcinoma cells they may also limit the impact of native anti-tumor immunity and that combining T cell-centric immunotherapy approaches (e.g. checkpoint blockade) with taxane agents requires additional scrutiny as these chemotherapies may in fact limit cytotoxic T cell migration into and through tumors.

**Figure 5.**
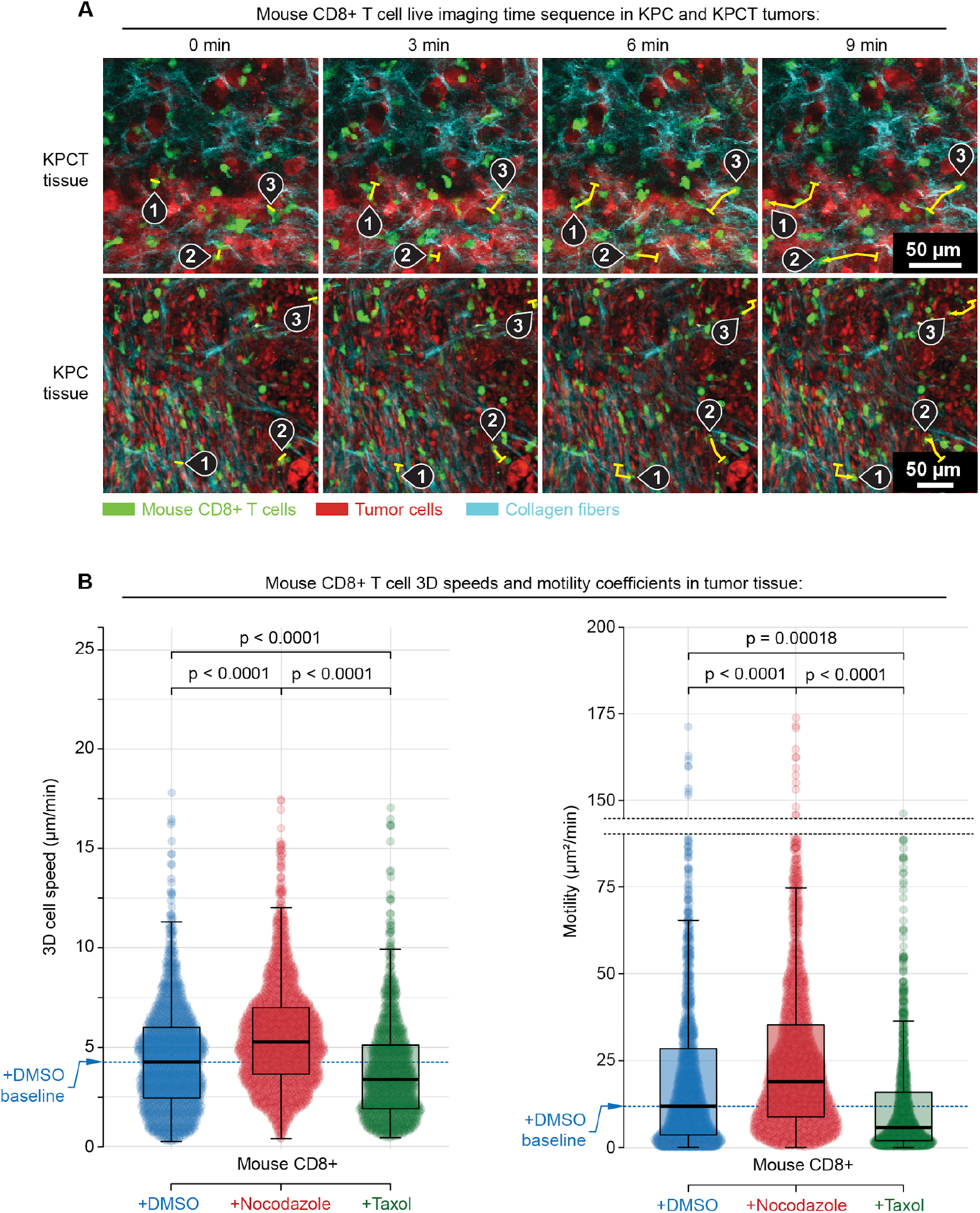
The microtubule-contractility axis regulates intratumoral migration of cytotoxic T cells. **(A)** Combined multiphoton excitation of fluorescence (MPE) and second harmonic generation (SHG) of primary murine CD8+ cytotoxic T cells (green) in paired live pancreatic tumor from *KPC* mice either expressing tdTomato specifically in carcinoma cells (*KPCT*; *top row*) or with all cells fluorescently labeled (*KPC; bottom row*). Sample migration tracks of individual T cells are highlighted using numerated pins along with their tracks (yellow curves; *start position - tick, end position - arrowhead*). Images are samples from imaging every 3 minutes for >1hr. **(B)** Speeds (*left*) and Motility Coefficients (*right*) for cytotoxic T cells migrating within 3D tumor microenvironments under control (+DMSO), MT destabilizing (+Nocodazole), and MT stabilized (+Taxol) treatment conditions, where MT destabilization leads to hypercontractile states and increased migration through stroma dense native pancreatic adenocarcinomas, while treatment with Taxol significantly decreases the ability of T cells to migration through native tumor microenvironments.

To further confirm our findings that the microtubule-contractility axis is a fundamental regulator of T cell migration in complex environments, and specifically test our hypothesis that GEF-H1 is a key mediator of microtubule to Rho-mediated contractility signaling, we employed CRISPR technology to knockout (KO) GEF-H1 in three independent primary human T cell lines (**Supplemental Figure 4**) and analyzed their 3D migration (**Figure 6**). In order to test our hypothesis that Rho activation from Nocodazole-induced MT instability is regulated by GEF-H1 we compared 3D migration of control Cas9 cells and GEF-H1 KO cells in the presence of Nocodazole (**Figure 6A**). Consistent with our hypothesis, under MT instability, migration is significantly decreased in GEF-H1 KO cells compared to control cells expressing Cas9, with overall motility cut in half (**Figure 6A-C**). However, remarkably, in separate experiments we discovered that knockout of GEF-H1 without MT destabilization significantly promotes migration (**Figure 6D-F**). In 3 primary activated human T cells lines, migration speed through 3D matrices is significantly increased (**Figure 6E**) while overall motility is significantly increased by more than 50% (**Figure 6F**), showing that genome engineering to eliminate GEF-H1 can result in engineered T cells that are more capable of migration throughout mechanically and structurally complex environments. Thus, overall these data demonstrates that approaches to alter T cell cytoskeletal-to-contractile machinery and signaling have the potential to significantly enhance T cell movement through complex solid tumor environments. Indeed, genome engineering strategies, or focused pharmalogic targeting, to alter the microtubule-contractility axis are likely to enhance T cell migration in the therapeutic setting where T cells are often not able to move throughout and sample an entire solid tumor volume.

**Figure 6.**
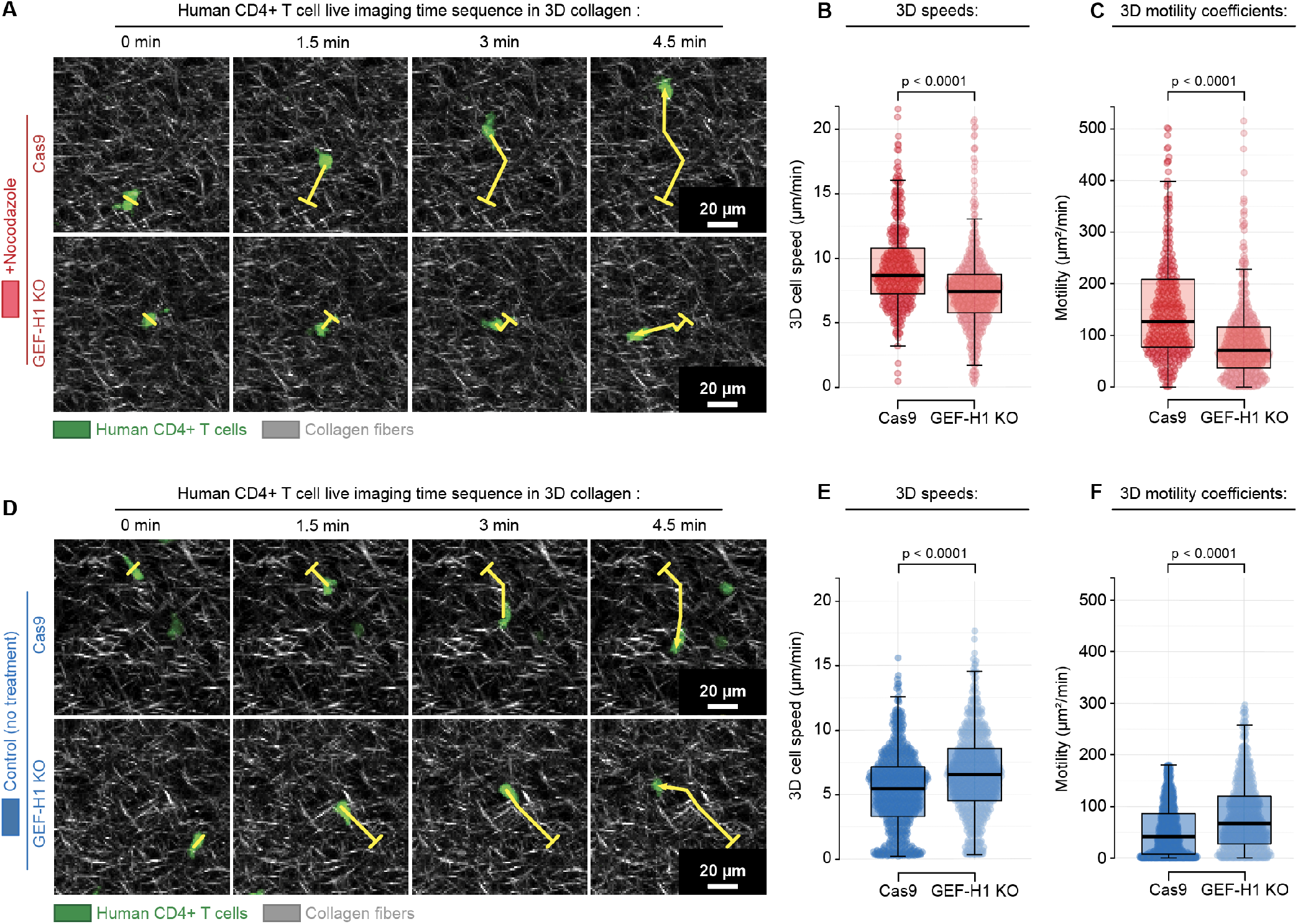
CRISPR GEF-H1 knockout (KO) prevents enhanced migration from microtubule destabilization but enhances migration of untreated T cells. **(A)** Combined multiphoton excitation of fluorescence (MPE) and second harmonic generation (SHG) of primary human T cells (green) migrating through 3D matrices. To test the hypothesis that increased contractility from destabilized microtubules is GEF-H1 dependent, we knocked out GEF-H1 using CRISPR and tested migration over time in the presence of destabilized MTs (+Nocodazole). Samples of migrating T cells are shown and tracks of individual cells are highlighted with their tracks (yellow curves; *start position - tick, end position - arrowhead*). Images are samples from imaging every 1.5 minutes for >1hr. **(B)** Speeds (*left*) and (**C)** Motility Coefficients (*right*) for control (Cas9) and GEF-H1 KO T cells migrating within 3D microenvironments under MT destabilizing conditions (+Nocodazole) showing that loss of GEF-H1 disrupts the contractility-enhanced migration from MT destabilization. **(D)** Combined MPE/SHG of control and GEF-H1 KO cells without destabilization of MTs. Samples of migrating T cells are shown and tracks of individual cells are highlighted using numerated pins along with their tracks (yellow curves; *start position - tick, end position - arrowhead*). (**E)** Speeds (*left*) and (**F)** Motility Coefficients (*right*) for control (Cas9) and GEF-H1 KO T cells migrating within 3D microenvironments with unperturbed MTs showing increased migration through complex 3D environments following genome engineering to knockout GEF-H1.

## DISCUSSION

The physical and molecular mechanistic principles regulating the dynamic interactions between migrating T cells and the 3D environment, e.g. the basis of T cell 3D locomotion throughout normal and solid tumor stroma, remain elusive, largely due to the overall complexity of 3D environments and undeciphered phenotype plasticity, such as coexisting amoeboid and mesenchymal modes of motility. Despite the widely acknowledged fact that the steric and structural density, architecture, and mechanics of solid tumors can prevent effective T cell infiltration or distribution (Elahi-Gedwillo et al., 2019; Hartmann et al., 2014; Kuczek et al., 2019; Salmon et al., 2012; Stromnes et al., 2017), the study of mechano- and structure-sensitive aspects of T cell motility has predominantly focused on flat 2D environments (i.e. more mesenchymal-like phenotypes) or confinement in microchannels (Jacobelli et al., 2010; Krummel et al., 2014; Saitakis et al., 2017; Tooley et al., 2009). In this study we attempted to comprehensively interlink the mechanistic and mechanosensing principles of 2D and 3D T cell motility. As a result, we principally advanced the concept of super-positioned amoeboid-mesenchymal phenotypes in T cells and deciphered a mechanosensing role during the dynamically adaptive balance between amoeboid-mesenchymal phenotypes that impacts 3D migration. By focusing on key controlling factors: microenvironment architecture, mechanics, and T cell myosin-driven contractility, we established that 3D amoeboid migration is proportional to actomyosin contractility, and provides a significant advantage with faster T cell motility in both “2.5D” and 3D environments. Importantly, the increase in T cell amoeboid behavior is inherently accompanied by increased nano-scale steric interactions with ECM texture, indicating the leading role of T cell shape shifting, with directed pseudopods, as a driver of T cells shifting to a more amoeboid locomotion. Furthermore, increasing T cell contractility either via direct RhoA activation or via the MT-GEF-H1 axis desensitized T cells to microenvironment mechanics, shifting T cells more toward an amoeboid mode, resulting in increased motility in dense 3D collagen-FN matrices and in *KPC* tumors. Thus, we define the starting basis for a new generation of T cell engineering strategies that address and tackle the issue of poor efficiency of T lymphocyte movement throughout architecturally and mechanically complicated and heterogeneous microenvironments.

Stromal cell populations and the ECM surrounding cancerous cells in the pancreas are believed to be critically involved in tumor growth, metastasis, and resistance to therapy (Feig et al., 2012; Provenzano et al., 2012), and also limit effective antitumor immunity from T cells due to immunosuppression and fibrotic barriers (Clark et al., 2007; Elahi-Gedwillo et al., 2019; Feig et al., 2013; Hartmann et al., 2014; Jiang et al., 2016; Kuczek et al., 2019; Stromnes et al., 2014). Yet, following specific strategies to alter the stroma, PDA becomes susceptible to immune checkpoint blockade (Feig et al., 2013; Jiang et al., 2016) and engineered T cells can overcome some of the physical and chemical barriers (Stromnes et al., 2015). Thus, therapies that rely on T cell function can be effective against PDA, and other solid tumors, in the right setting. Yet, one key obstacle to maximally effective T cells therapies in solid tumors is their inability to efficiently navigate through heterogeneous tumor microenvironments to sample the entire tumor volume within a functional time domain. Here we present an advance to overcoming this obstacle and show that perturbing the microtubule-to-contractility axis, and in particular activation of Rho-mediated contractility, results in T cells with increased capacity to move through solid tumor microenvironments. Thus specific strategies, such as pharmacologic interventions and genome engineering, to enhance contractility either through elements of adhesion-to-cytoskeleton-to-Rho signaling pathways or direct alteration of contractility-governing molecular motors have a role in T cell engineering to generate “mechanically optimized” therapeutic cells. This first volley to overcome barriers in the tumor microenvironment provides a foundation from which to build upon. With the recent advances in genome engineering, such as the use of CRISPR technology here, we are well poised for future studies to test rational engineering strategies to improve the ability of T cells to move throughout and sample mechanically and chemically complex tumor microenvironments in order to develop next-generation immune-focused therapies to improve patient outcomes.

## MATERIALS and METHODS

### RESOURCES TABLE

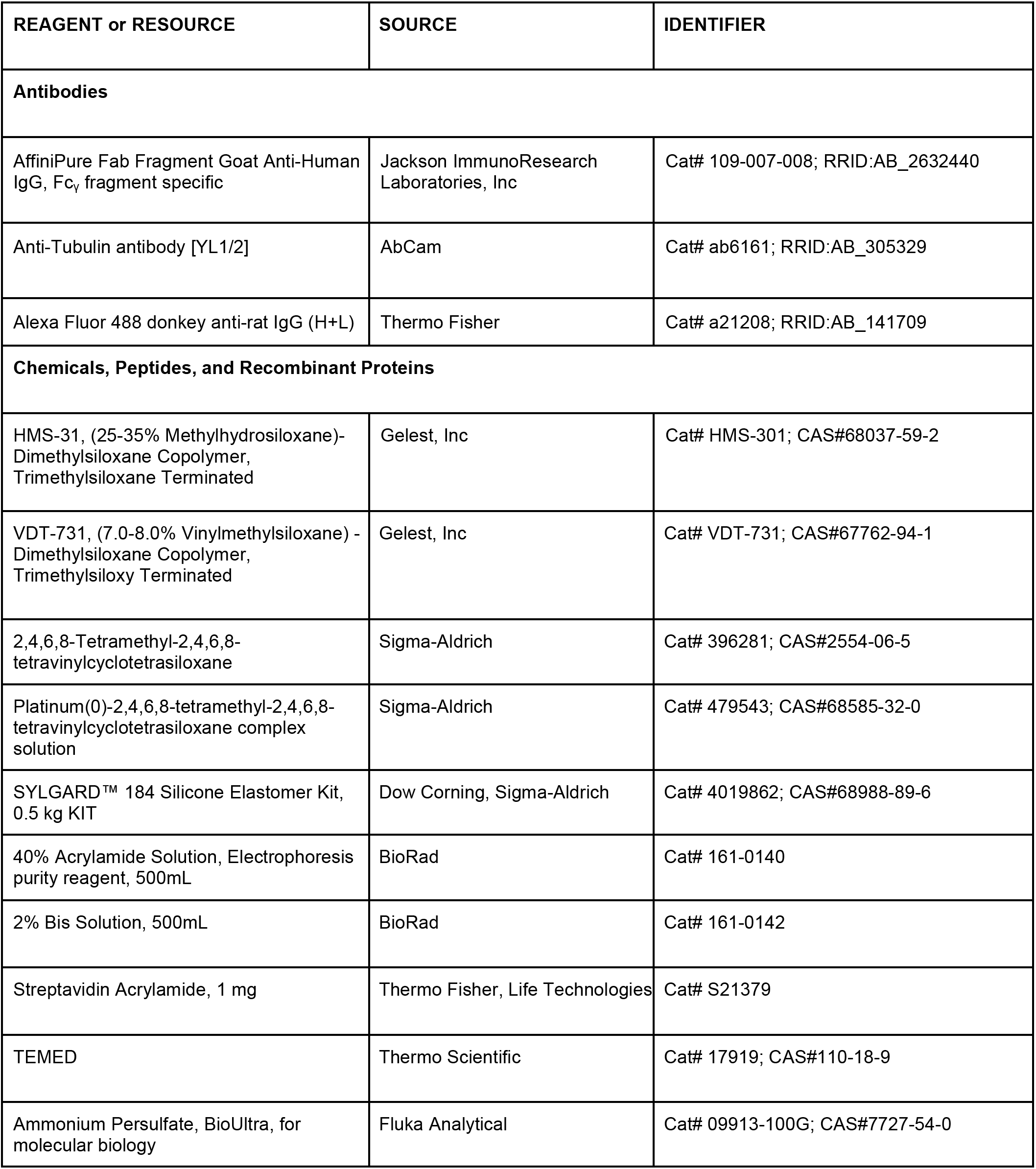

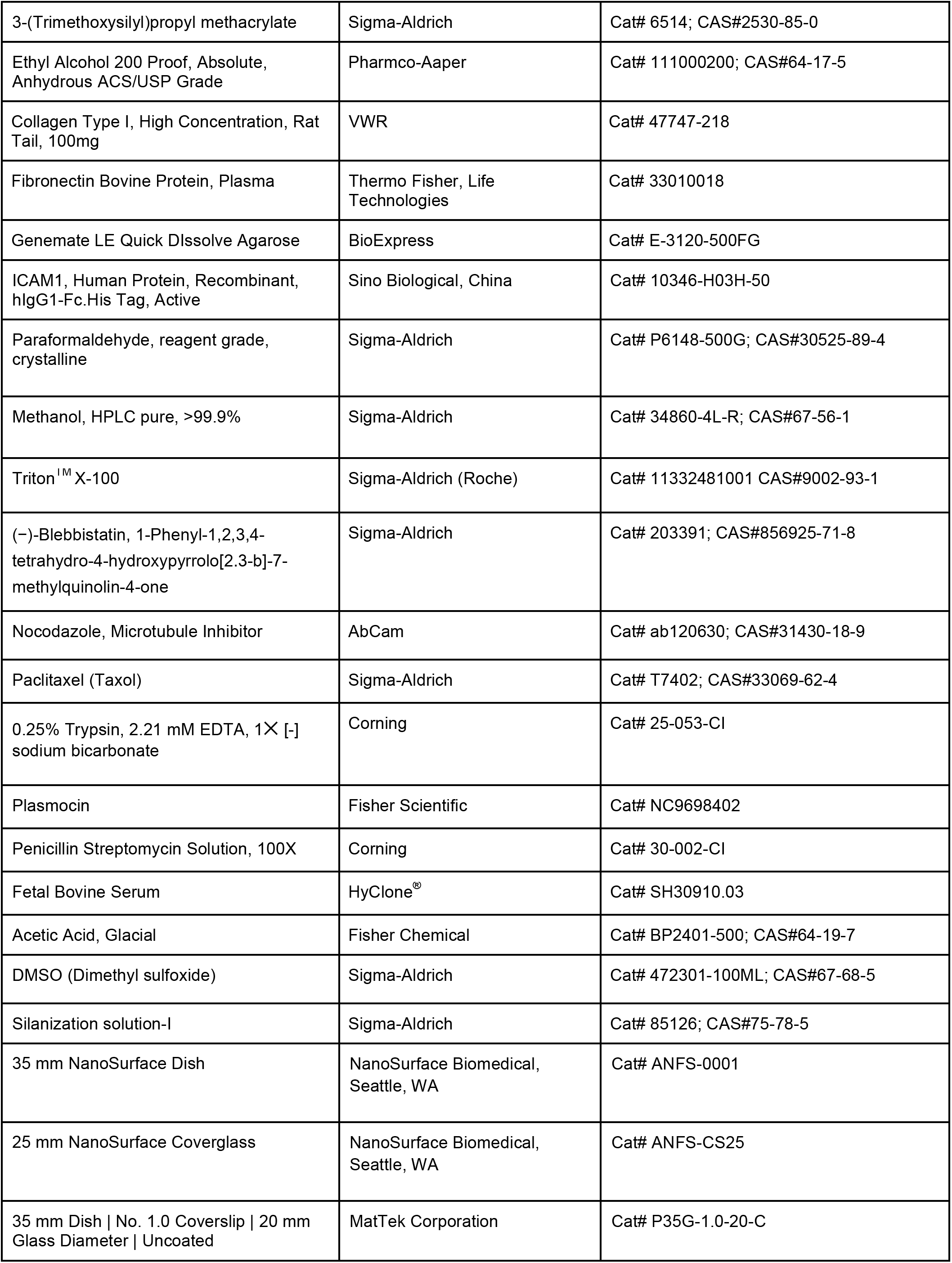

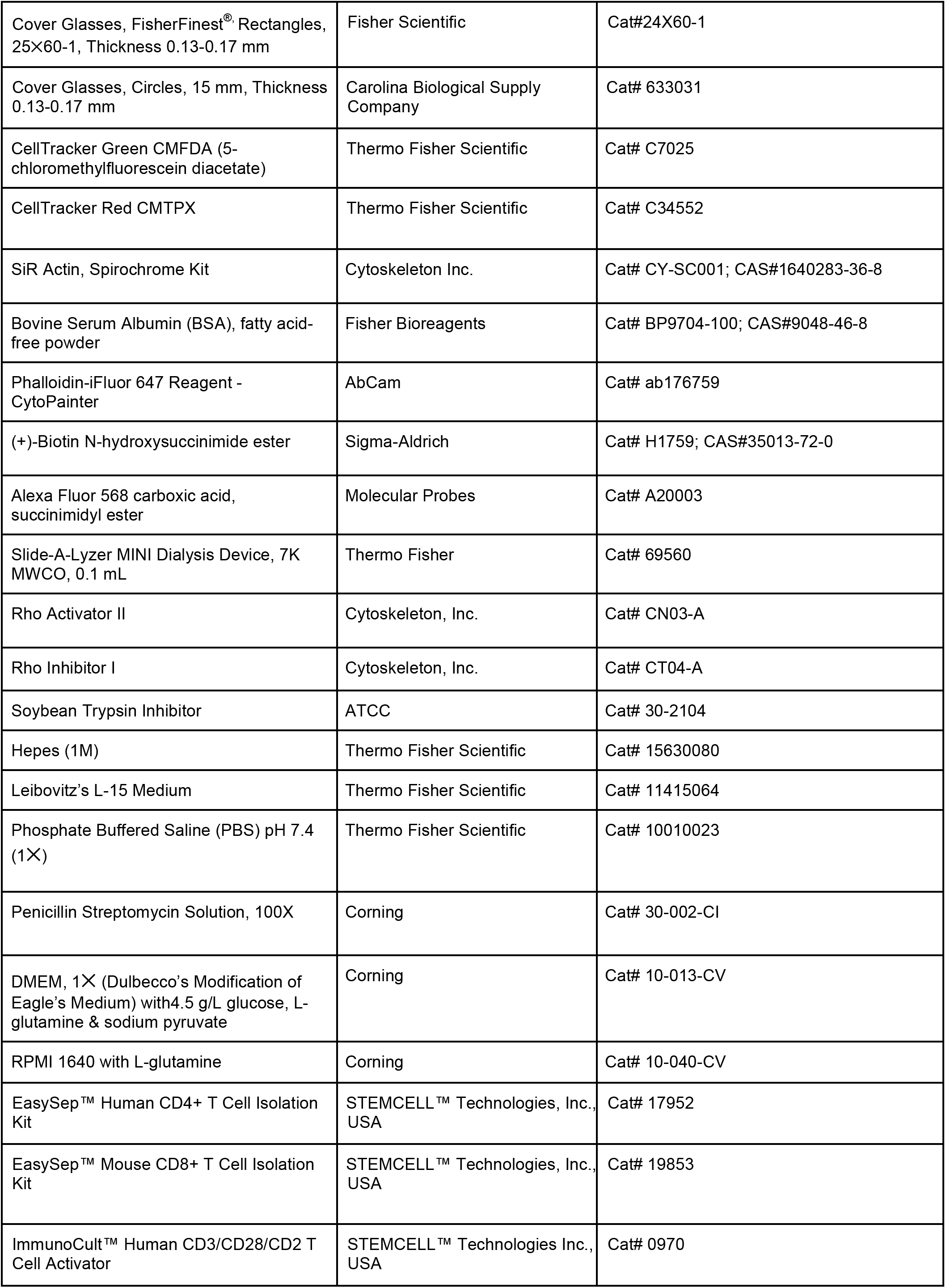

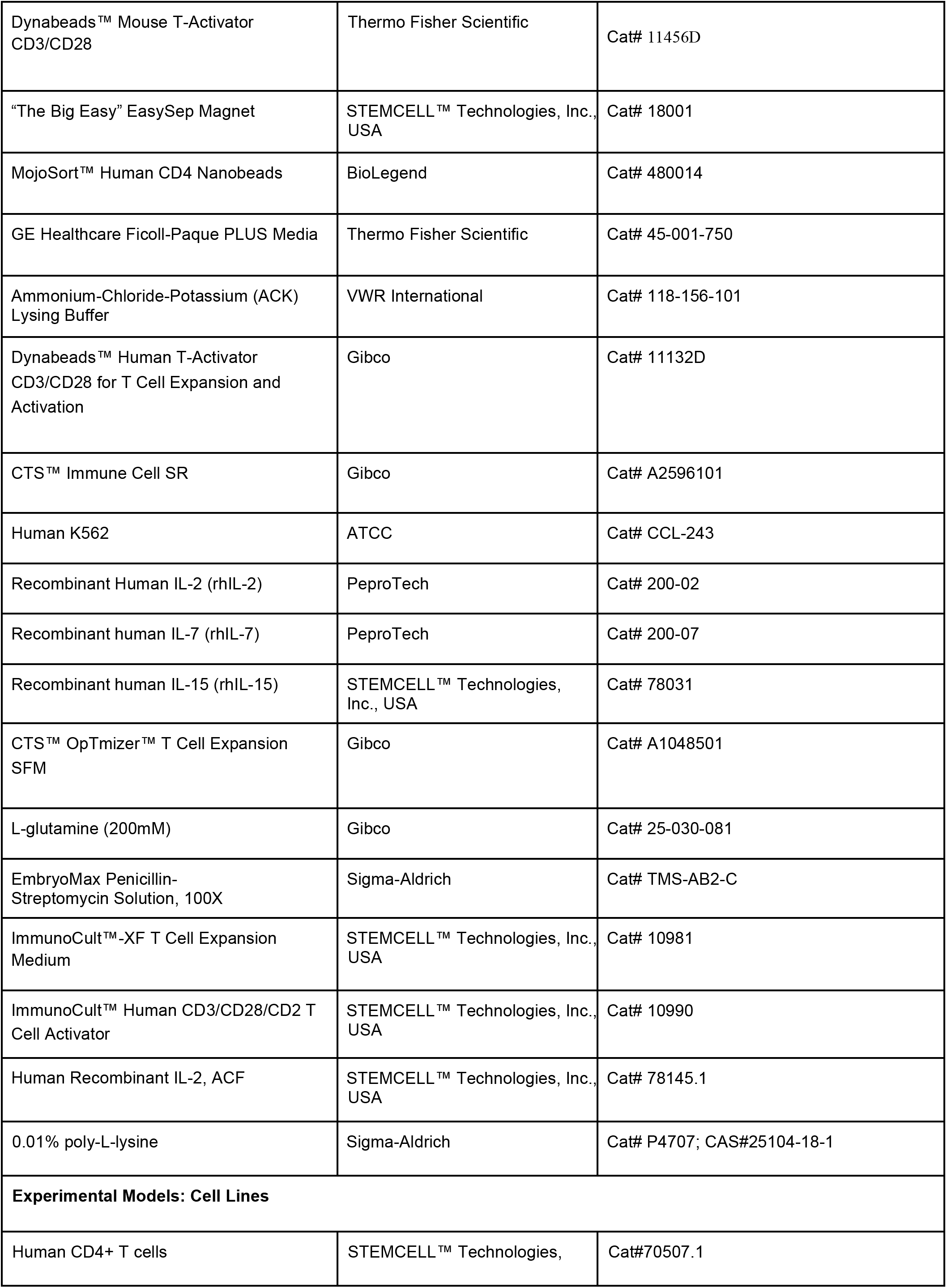

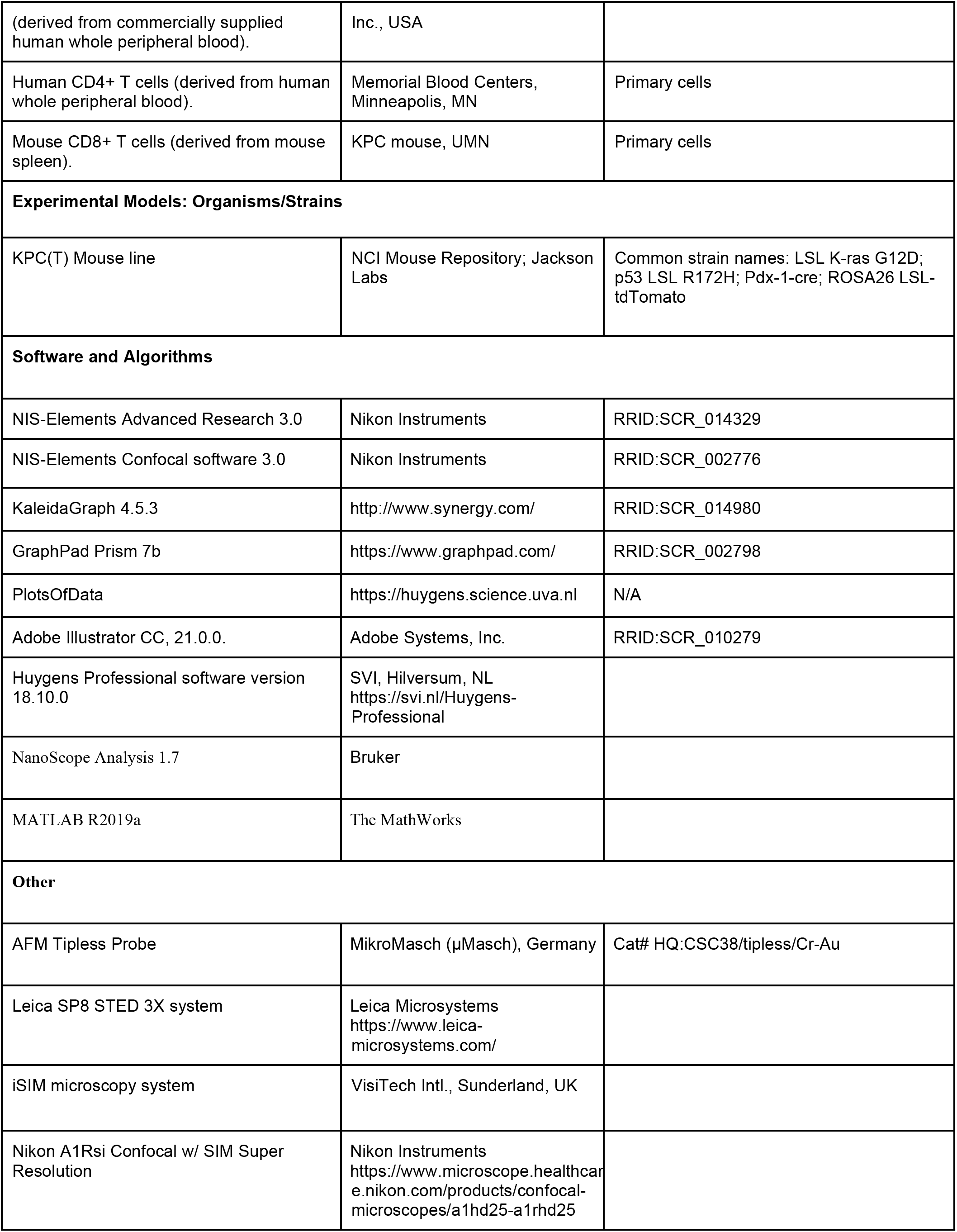

### Cell culture

Human CD4+ T cells were produced by CD4+ cell isolation and purification from the whole human blood, supplied by STEMCELL™ Technologies Inc., where blood was tested and negative for Hepatitis C and HIV. CD4 T cell isolation and purification was performed with EasySep™ Human CD4+ T Cell Isolation Kit (STEMCELL™ Technologies Inc., USA) and then T cells were cultured, activated and expanded in ImmunoCult™-XF T Cell Expansion Medium (STEMCELL™ Technologies Inc., USA) with the addition of ImmunoCult™ Human CD3/CD28/CD2 T Cell Activator and Human Recombinant Interleukin 2 (IL-2, STEMCELL™ Technologies Inc., USA) as per STEMCELL™ Technologies Inc. commercial protocol. For CRISPR protocols, human CD4+ (hCD4+) cells derived from human whole peripheral blood, were activated and expanded using ImmunoCult™ Human CD3/CD28/CD2 T Cell Activator (STEMCELL™ Technologies Inc., USA), following the manufacturer’s recommendation, in ImmunoCult™-XF T Cell Expansion Medium (STEMCELL™ Technologies Inc., USA) supplemented with Human Recombinant IL-2 (STEMCELL™ Technologies Inc., USA). For CRISPR experiments, peripheral blood mononuclear cells (PBMCs) from de-identified healthy human donors were obtained using Trima Accel leukoreduction system (LRS) (Memorial Blood Centers, Minneapolis, MN) and further purified using ammonium chloride-based red blood cell lysis and a Ficoll-Paque gradient. CD4+ T cells were isolated from the PBMC population by immunomagnetic positive selection using the MojoSort Human CD4 Nanobeads (BioLegend, San Diego, CA) and an EasySep Magnet (STEMCELL™ Technologies Inc., USA). Mouse CD8+ T (mCD8+) were isolated from tumor bearing *KPC* or *KPCT* mice (*Kras^G12D/+^;p53^R172H/+^;Pdx1-Cre* or *Kras^G12D/+^;p53^R172H/+^;Pdx1-Cre;ROSA^Tdtomato/+^*, respectively), using EasySep™ Mouse CD8+ T Cell Isolation Kit (STEMCELL™ Technologies Inc., USA) following the manufacturer’s recommendations. After isolation, mCD8+ T cell activation and expansion was performed using Dynabeads Mouse T-Activator CD3/CD28 (Thermo Fisher Scientific), following the manufacturer’s recommendations in ImmunoCult™-XF T Cell Expansion Medium supplemented with Human Recombinant IL-2 for at least 4 days. Magnetic separation was done to separate beads from cells. Cells can be used immediately or can be frozen down in freezing medium (15% DMSO in FBS) for future use. Note that frozen T cells were given an incubation/recovery period (*t*≥24 hours) in ImmunoCult™ expansion medium, supplied with human IL-2 after cell thawing, in order to avoid the effects of cold exposure shock. All cell work was approved by the University of Minnesota Institutional Biosafety Committee and followed institutional and NIH guidelines.

### High precision nano-stamps

A more detailed, step-by-step protocol for this procedure is described elsewhere (Tabdanov, Zhovmer et al., 2020). Fabrication of elastic ICAM1 or FN nanopatterns is a challenging task due to the van-der-waals and capillary interactions between the nano-stamp and the printed surface that provoke a collapse of the soft PDMS nano-stamps onto the printed intermediate (glass) surface. To address these issues and achieve high precision of nanopatterns on elastic platforms we substituted regular PDMS nano-stamps with composite stamps, veneered with a submillimeter-thick hard PDMS (hPDMS) for non-collapsing high-definition printing surfaces (Schmid and Michel, 2000; Tabdanov et al., 2018b). For the hPDMS preparation protocol, please see “*hPDMS formulation*” section. In order to cast the nano-printing surface, we used commercially manufactured polyurethane nano-surfaces as the casting matrices (NanoSurface Biomedical, Seattle, WA). Clean textured nano-surface (NanoSurface Biomedical, Seattle, WA) disks were glued onto the glass platform with SuperGlue® (Loctite, USA), silanized with silanizing solution-I as per the commercial protocol (Sigma Aldrich), coated with ≤0.5mm hPDMS by gentle spreading with soft Parafilm-made spatula (Hach, USA), cured at 70°C for 30 minutes and subsequently cast with regular PDMS to the layer final thickness of 8 mm (rPDMS; 1:5 curing agent/base ratio, Sylgard-184, Dow Corning). Cured (at 70°C for ~1 hour) composite nano-stamps were peeled, and cut into 5×5 mm or 1×1 cm pieces and used as the ready-to-use nano-stamps. To fabricate ICAM1 elastic nanotectures the Fab anti-Fc antibody fragment protein (Jackson Immunoresearch, USA) was prelabeled with a fluorescent tag and a biotin group to ensure both its fluorescent visibility in nanopatterns and cross-linking to the streptavidin-functionalized PAA gels, respectively. Briefly, 20 μL of 1 mg/mL antibody sample was incubated for 1 hour with 5 μL of ((+)-biotin N-hydroxysuccinimide ester, Sigma-Aldrich; as per the commercial protocol) and 5 μL of fluorescent tag kit (Alexa Fluor® succinimidyl esters, Invitrogen, Molecular Probes®; as per the commercial protocol). Labeled protein then was dialysed overnight in Slide-A-Lyzer MINI Dialysis Device, 7K MWCO (Thermo Fisher) overnight at 4°C in cold PBS, then stored at 4°C in the darkness. 10 μL droplets of 0.1 mg/mL labeled antibody solution were then placed atop of the 5×5 mm or 1×1 cm square micro- or nano-stamps. To ensure a proper coverage and effective stamp surface coating with labeled Fab anti-Fc antibody fragment protein, the protein solution droplet was “sandwiched” between the stamp’s printing surface and 15 mm round glass coverslip (Carolina, USA), which had been baked in the furnace for 5-10 hours at 450°C.

### High precision 2D nanocontact printing

A detailed step-by-step protocol for this procedure is described elsewhere (Tabdanov and Zhovmer et al., 2019). Briefly, using the nano-stamps, we first printed Fab anti-Fc antibody fragment protein nanopatterns onto the “intermediate” glass surface (Tang et al., 2012), which then were cross-linked to polymerizing PAA gels premixes by their biotin tags to streptavidin-conjugated polyacrylamide (Streptavidin-acrylamide, Thermo Fisher). For that 7-10 μL of PAA premix of desired G’ was polymerized in the “sandwich” fashion between an “intermediate” patterned surface and glass-bottom 35 mm Petri dishes (MatTek Corp., Ashland, MA), activated with 3-(trimethoxysilyl)propyl methacrylate (Sigma-Aldrich) in ethyl alcohol (Pharmco-Aaper) and acetic acid (Fisher Chemical) as per the commercial protocol. For specific G’ value formulations, please see the “*PAA elastic gels premixes*” section. 3-(trimethoxysilyl)propyl methacrylate-functionalized glass surface establishes covalent bonds with the PAA gel upon its curing. Polymerized PAA “sandwiches” then were subjected to hypotonic reversible swelling in deionized water (overnight) for a gentle coverglass release from PAA gel. The resultant fluorescent PAA-nanopatterns of Fab anti-Fc antibody fragment protein were incubated overnight with 1 mg/mL ICAM1-Fc chimeric protein (Sino Biological, China) in cold PBS (Gibco, USA) at 4°C, rinsed, and used for experiments.

### Fabrication of elastic “2.5D” nanotextures

A detailed step-by-step protocol for this procedure is described elsewhere (Tabdanov and Zhovmer et al., 2019). Briefly, nanotextures were cast from PAA gel premixes of chosen shear modulus (G’). For specific G’ value formulations, please see the “*PAA elastic gels premixes*” section. As the nanotexture casting master we used polyurethane-based texturized nanosurfaces (NanoSurface Biomedical, Seattle, WA), which was used to manufacture hPDMS-based molds (please see the hPDMS making protocol in “*hPDMS formulation*” section). 5-7 μL droplet of freshly prepared liquid hPDMS premix was sandwiched between polyurethane-based texturized nanosurfaces and a clean activated (baked at 450°C for 12 hours, then ozone or regular air plasma-treated for 5 minute) coverglass slide. hPDMS droplet is allowed to spread into thin layer (5-10 minutes at ~20°C), then baked at 70°C for 1 hour. Cured hPDMS “sandwich” is then separated from the molding surface by manual peeling of the hPDMS+glass part of the “sandwich” from the casting polyurethane surface. Peeling process releases hPDMS surface from polyurethane casting master, leaving hPDMS layer attached to the activated glass. The resultant molding nanosurface is then cut in 1×1 cm squares by diamond pencil scribbling (on the reverse side of nano-surface) and precoated with biotinylated and fluorescent tag-labeled anti-Fc Fab antibody fragment (Jackson Immunoresearch; 0.1 mg/mL PBS solution, 4°C, in wet chamber overnight). Alternatively, for fabrication of fibronectin-functionalized elastic nanotextures, 0.1 mg/mL fibronectin protein (FN) PBS solutions (Bovine plasma fibronectin, Thermo Fisher, Life Science, USA) was used, labelled following the same protocol, as for Fab anti-Fc protein (see section “High precision nano-stamps”). After the incubation with the protein solution, the molding nanosurface was gently rinsed in deionized water, blow dried under a filtered air, argon or nitrogen jet. Streptavidin-conjugated polyacrylamide premix of volumes not greater than 0.5 mL was degassed in a vacuum chamber or in an ultrasonication water bath for 1 hour. In order to prevent TEMED evaporation during the procedure, TEMED is added after the degassing session. 7-10 μL droplet of PAA premix of desired G’ value was polymerized in the ‘‘sandwich’’ fashion between protein-coated nanosurface and glass-bottom 35 mm Petri dishes (MatTek Corp., Ashland, MA), activated with 3-(trimethoxysilyl)propyl methacrylate (Sigma-Aldrich) in ethyl alcohol (Pharmco-Aaper) and acetic acid (Fisher Chemical) in a vacuum chamber. After PAA curing the resultant textured patterned elastic chip was placed overnight into cold (i.e. at 4°C) deionized water for PAA reversible hypotonic ‘‘swelling.’’ Then the casting surface was gently peeled from the polymerized PAA surface. For a better release of the sterically interactive nano-mold, hypotonically treated PAA ‘‘sandwiches’’ were optionally ultrasonicated in the water bath for 10 sec. Released FN elastic nanosurface is ready for use immediately. Prepared elastic Fab anti-Fc-functionalized PAA nanotextures were incubated with 20 μg/mL human ICAM1-Fc chimeric protein (Sino Biological, China) in cold PBS (4°C, 12 hours), rinsed in PBS three times, and used for the T cell adhesion and contact guidance assays.

### hPDMS formulation

For hard PDMS (hPDMS) we mixed 3.4 g of VDT-731 (Gelest, Inc.), 18 μL of Pt catalyst (Platinum(0)-2,4,6,8-tetramethyl-2,4,6,8-tetravinylcyclotetrasiloxane complex solution) (Sigma-Aldrich) and one drop of cross-linking modulator (2,4,6,8-Tetramethyl-2,4,6,8-tetravinylcyclotetrasiloxane) (Sigma-Aldrich). Next, immediately before use, we added 1 g of HMS-301 (Gelest, Inc.) and thoroughly mixed it for 30 sec on vortex mixer (Odom et al., 2002; Schmid and Michel, 2000).

### PAA elastic gels premixes

We chose to control PAA mechanical rigidity via modulation of concentration for both 40% acrylamide (40% AA) base (BioRad) and its cross-linking molecular chain, 2% bis-AA (BioRad) as described elsewhere (Fischer et al., 2012; Plotnikov et al., 2014). Additionally, streptavidin-acrylamide (Thermo Fisher) was added to the final concentration of 0.133 mg/mL to enable PAA gels cross-linking with biotinylated proteins of interest. Briefly, for preparation of 50 μL of G’ = 2.3 and 50 kPa PAA gel premixes, respectively, the following components were mixed: 40% AA: 9.33 and 15 μL; 2% bis-AA: 1.88 and 14.40 μL; 2 mg/mL streptavidin-AA: 3.33 and 3.33 μL; 10X PBS: 5 and 5 μL; deionized milli-Q water: 30 and 11.17 μL; TEMED: 0.1 and 0.1 μL; 10% APS: 1 and 1 μL. The premix solutions were degassed and stored at 4°C before use.

### T cell contact guidance and migration analysis on “2.5D” platforms

Cell tracking on the 2D and “2.5D” surfaces was performed with the TrackMate plug-in in ImageJ. T cells were imaged with differential interference contrast (DIC) microscopy at 10 sec intervals using a 20x air objective (Nikon Instruments, Japan) and tracking was also confirmed manually, one cell at a time, at each interval. Cells were maintained at 37°C in 5% CO_2_ (Tokai Hit, Japan) for the duration of live cell imaging session. Images were captured at resolutions of 512×512 or 1024×1024 pixels.

### Super-resolution imaging and quantification of human T cells invasiveness on nanoarchitectures

Super-Resolution Stimulated Emission Detection (STED) microscopy (Figure 1A, Figure S1, Figure 4A) was performed using a commercial Leica SP8 STED 3× system (Leica Microsystems, Mannheim, Germany), equipped with a white light laser with continuous spectral output between the wavelengths of 470 nm to 670 nm, and a 592 nm, 660 nm and a pulsed 775 nm STED depletion lasers, to obtain time-gated STED images on 3 hybrid detectors. Given the complexity and varying depth of the sample, we have used the STED WHITE Glycerin objective lens (HC PL APO 93×/1.30 GLYC motCORR) (Leica Microsystems) which is advantageous for depth imaging due to the motorized correction collar allowing precise and swift adjustment of optical lenses to specimen inhomogeneity. SiR-actin labeled samples (per commercial protocol) placed in 35 mm culture dishes with number 1.5 cover glass bottom (MatTek Corporation, Ashland, MA) containing 250 μl glycerol (90%) in PBS were imaged in sequentially as follows: first sequence STED for SiR-actin (via 647 nm excitation and 660-730 nm emission range) on gated (0.7-6.5 ns time gating) hybrid detector using 775 nm (25% power) as STED depletion laser for best lateral resolution; Second and third sequences were confocal settings for Hoechst and Alexa488 (labeling nucleus and surface of the gel) respectively, via two sequential excitations (405 nm, and 488 nm) and two emission ranges (410-465 nm, and 495-555 nm) respectively, on gated (0.3-6.5 ns) hybrid detectors. Imaging was performed with a scan speed of 600 lines per second, scanning bidirectionally, a pixel size of 30-35 nm (1024×1024 pixels), and 6-line averages, pinhole of 0.7 Airy units and Z-stacks were collected at 0.140 μm-depth intervals throughout the depth of the sample. We deconvoluted images using Huygens Professional software version 18.10.0 (SVI, Hilversum, NL) with the classical maximum likelihood estimation algorithm. We then inspected and reconstructed 3D data using Clear Volume plugin (FIJI). Still frames were saved and montaged using Adobe Photoshop CC.

Data collection for quantification of cell invasiveness (Figure 1E) was performed using the instant structured illumination microscopy (iSIM) by an Olympus IX-81 microscope (Olympus Corp., Tokyo, Japan) equipped with an Olympus UPLAPO-HR 100x/1.5 NA objective, two Flash-4 scientific CMOS cameras (Hamamatsu Corp. Tokyo, Japan), an iSIM scan head (VisiTech Intl., Sunderland, UK), and a Nano-Drive piezo Z stage (Mad City Labs, Madison, WI). The iSIM scan head included the VT-Ingwaz optical de-striping unit (VisiTech Intl., Sunderland, UK). Image acquisition and system control was through MetaMorph Premiere software (Molecular Devices, LLC, San Jose, CA). Images were deconvolved using the specific for iSIM commercial plugin from Microvolution (Cupertino, CA) in FIJI.

### Atomic force microscopy

Human CD4+ T cells were plated on a glass-bottom dish (Willco Wells) pre-coated with either ICAM-1 (Life Technologies) or 0.01% poly-L-lysine (Sigma-Aldrich) and immersed in culture media solution (Life Technologies). Force spectroscopy atomic force microscopy experiments were performed using a Bruker Bioscope Catalyst AFM system (Bruker) mounted on an inverted Axiovert 200M microscope (Zeiss) equipped with a confocal laser scanning microscope 510 Meta (Zeiss) and a 40x objective lens (0.95 NA, Plan-Apochromat, Zeiss). The hybrid microscope instrument was placed on an acoustic isolation table (Kinetic Systems). During AFM experiments, T cells were maintained at physiological relevant temperature 37°C using a heated stage (Bruker). A soft silicon nitride tipless AFM probe (HQ:CSC38/tipless/Cr-Au, MikroMasch) was used for T cell’s compression. The AFM microcantilevers were pre-calibrated using the standard thermal noise fluctuations method. The estimated spring constants for microcantilevers used were 0.07 – 0.1 N/m. After calibration, the AFM probe was moved on top of a rounded T cell. Five to ten successive force curves were performed on each T cell. The deflection set-point was set to 20 nm yielding applied forces between 1.5 to 2 nN.

### Analysis of AFM data: Determination of T cell’s cellular surface tension

All AFM force-distance curves measurements were analyzed using a custom-built MATLAB (The MathWorks) code to calculate the cellular surface tension. For curves fitting, indentation depths between 0-400 nm were relatively consistent in yielding good fits (R^2^>0.9). Curves with poor fits R^2^<0.9 were discarded from the analysis. Additionally, we discarded noisy force curves and/or curves that presented jumps possibly due to cantilever and plasma membrane adhesion, slippage, or very weakly adhered and moving cells. T cell surface tension (*T*; pN/μm) was computed by fitting each recorded force-distance curve with the surface tension formulae described in (Cartagena-Rivera et al., 2016; Logue et al., 2015) that defines the force balance relating the applied cantilever force with the pressure excess inside the rounded cells and the corresponding surface tension; 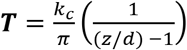, where *T* is the cellular surface tension, *k_c_* is the AFM cantilever spring constant, *Z* is the Z-piezo extension, and *d* is the cantilever mean deflection. Additionally, the T cells intracellular hydrostatic pressure (*P*; Pa) was calculated by using Laplace’s law for spheres; 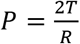, where *P* is the intracellular hydrostatic pressure and *R* is the initial radius of the T cell.

### T cell genome engineering with CRISPR

Guide RNAs (sgRNAs) were designed to hARHGEF2 using the Synthego CRISPR design tool (https://design.synthego.com/). The top two recommended guides were obtained as modified sgRNAs (Synthego, Menlo Park, CA). Both sgRNAs were screened in K562 cells and the most efficient gRNA (Sequence: 5’-GAGGUGCCCAUUGGUAUAGC-3’) based on knockout score was used in subsequent experiments with primary human T cells. For T cell culture, T cells were maintained in OpTmizer CTS T cell Expansion serum-free media (SFM) containing 2.5% CTS Immune Cell Serum Replacement (SR) (ThermoFisher, Waltham, MA), L-Glutamine, Penicillin-Streptomycin, N-Acetyl-L-cysteine (10 mM), rhIL-2 (300 IU/mL), rhIL-7 (5 ng/mL), and rhIL-15 (5 ng/mL), at 37°C with 5% CO_2_. T cells were activated with Dynabeads Human T-Activator CD3/CD28 (ThermoFisher, Waltham, MA) at a 2:1 bead:cell ratio for 48 h prior to electroporation. T cells were maintained at ~1×10^6^/mL in normal tissue culture flasks for experiments optimizing editing efficiency. For T cell electroporation, after 48 hours, Dynabeads were magnetically removed and cells were washed with PBS once prior to resuspension in appropriate electroporation buffer. 1×10^6^ Primary Human T cells were electroporated using the 4D-nucleofector (Lonza, Basel, Switzerland) and a P3 Primary Cell 4D-Nucleofector™ X Kit (V4XP-3032). 1.5 μg CleanCap Cas9 mRNA (TriLink Biotechnologies, San Diego, CA) and 1 μg of modified gRNA were added to 1×10^6^ cells in 20 μL of the recommended electroporation buffer. The mixture was electroporated using the program EO-115. Cas9 mRNA alone was used as a control for all conditions. Following electroporation, T cells were allowed to recover in antibiotic-free medium at 37 °C, 5% CO2 for 20 min, then cultured in complete CTS OpTmizer T cell Expansion SFM as described above. Genomic DNA was taken from T cells 7 days post electroporation by spin column-based purification. Cas9 efficiency was analyzed on the genomic level by PCR amplification of CRISPR-targeted loci (Forward sequence: 5’-AGGGAGATGAGTGGCAACAG-3’; Reverse sequence: 5’-CAGCTGGGGATCAGAGAGAA-3’), Sanger sequencing of the PCR amplicons (Eurofins Genomics, Louisville, KY), and subsequent analysis of the Sanger sequencing traces using the ICE web app developed by Synthego (https://ice.synthego.com/).

### Preparation of 3D collagen matrices

Collagen matrices preparation was adapted from (Ray et al., 2017b) with modifications. Briefly, high concentration rat-tail collagen I (VWR) was neutralized with a 1:1 ratio of 100 mM HEPES (Thermo Fisher Scientific) in 2X PBS and completed to a final concentration of 3 mg/mL with DMEM (Corning) supplemented with 10% FBS (which introduces FN). The mixture was allowed to sit in ice for 5 min, after which 350 mL was pipetted into a standard 24-well plate. The gels were left polymerizing for 20 min, in the hood, at room temperature and then incubated at 37°C with 5% CO_2_ for 3 h and subsequently overlaid with DMEM (Corning) and incubated overnight at 37°C with 5% CO_2_. For some instances of linear microscopy analysis, we imaged the collagen matrix via fluorescence instead of second harmonic generation imaging. To fluorescently label collagen-I matrix, we added 1:20-1:100 (volume) of 1 mg/ml fluorescent tag (Alexa Fluor® succinimidyl esters of the desired excitation/emission spectrum, Invitrogen™, Molecular Probes®) to the gel premix immediately before polymerization.

### T cell migration in 3D collagen matrices

3×10^5^ viable hCD4+ or mCD8+ T cells were stained with 1 μM of CellTracker Green CMFDA (5-chloromethylfluorescein diacetate; Thermo Fisher) for 5 minutes, protected from light, at 37°C. Lymphocytes were centrifuged at 300 g for 5 min and washed twice in L-15 plus 1% FBS and resuspended in the same media to a volume of 200 μL. Excess overlaid media from the collagen matrices was removed completely prior to seeding T-cells. The 3×10^5^ T cells in 200 μL were gently pipetted on top of the collagen matrix, gently swirled to evenly distribute the cell suspension and incubated for 30 min at 37°C and 0% CO_2_ to allow lymphocytes to infiltrate the gel matrix. After incubation, the remaining media on top of the matrix was removed and washed once to remove cells that did not infiltrate/adhere to the gel. Gel was carefully detaching from all sides using a pipette tip and transferred to a 35 mm dish using tweezers. A slice anchor (Warner Instruments) was used to hold the gel in place and was overlaid with 5 mL of L-15 media plus 1% FBS in control (DMSO), 70 nM Taxol, 10 μM Nocodazole, 5 μg/mL RhoA Activator II (Cytoskeleton, Inc.) or 2 μg/mL Rho A Inhibitor I (Cytoskeleton, Inc.) conditions and imaged immediately after designated incubation time as per commercial protocol (Cytoskeleton Inc.). Briefly, for Rho A activation or inhibition, prior to staining T cells with CMFDA, T cells were resuspended in serum free RPMI with 5 μg/mL and 2 μg/mL of activator and inhibitor for 3 and 4 hours, respectively.

### Mouse tumor slice culture

Genetically engineered *Kras^G12D/+^;p53^R172H/+^;Pdx1-Cre* (*KPC*) and fluorescent reporter *KPC* mice [*Kras^G12D/+^;p53^R172H/+^;Pdx1-Cre;ROSA^Tdtomato/+^* (*KPCT*)] were used as a faithful mouse model for pancreatic cancer (Elahi-Gedwillo et al., 2019; Hingorani et al., 2005; Provenzano et al., 2012) as approved by the Institutional Animal Care and Use Committee of the University of Minnesota. Freshly explanted *KPC* and *KPCT* tumors were harvested at endpoint and placed in sterile ice cold sterile PBS with soybean trypsin inhibitor (STI; ATCC). 1.5% agarose (Genemate) gel was prepared and super glued (Loctite, USA) onto the vibratome (Vibratome Company) cutting stage to create a support for the tumor which was super glued directly in front of the agarose gel. The stage was overlaid with ice cold PBS. The sectioning parameters were: speed 3 mm/s amplitude 8 mm and 350 μm thickness. Slices were placed in ice cold sterile PBS with STI for transport and subsequently cultured (4 days maximum), as described previously (Jiang et al., 2019) with modifications. Briefly, multiple slices were transferred, using tweezers, and placed flat on 0.4 μm, 30mm diameter cell culture inserts (Millipore Sigma) pre-coated with collagen gel mixture, as described above but completed with 1× PBS instead of growth media, in a 6-well plate, with RPMI 1640 supplemented with 10% FBS, 1% penicillin and streptomycin, 14.5 mM Hepes, 5 μg/mL plasmocin (Invivogen) and 10 μg/mL of STI in a 37°C humidified incubator at 5% CO_2_. Single slices can be cultured similarly in a 0.4 μm, 12 mm diameter cell culture inserts (Millipore Sigma) in a 24-well plate. The culture media was changed daily.

### T cell migration in tumor slices

*KPC* slices were transferred to a 24 well plate and cultured in L-15 media supplemented with 10% FBS, 1% penicillin and streptomycin, 5 μg/mL plasmocin and 10 μg/mL of STI with 5 μM of CellTracker Red CMTPX (Invitrogen) for 15-20 minutes at 37°C with 0% CO_2_ incubator. *KPCT* slices were not stained with CellTracker Red since carcinoma cells can be visualized via red fluorescence. Slices were washed twice in the cell tracker free media. Individual slices were then transferred using tweezers to the 12 mm culture inserts in the 24-well plate. 1-2×10^5^ mCD8+ T cells were stained, as described above, and concentrated in 100 μL of L-15 media supplemented with 10% FBS and immediately added onto the slice, in the inserts, as to concentrate the cells on the slice and let the cells adhere/migrate on the tissue for 1 hr at 37°C with 0% CO_2_ incubator. Slices were then handled with tweezers and gently washed in the same media to remove excess T cells that did not migrate/adhered into the tissue and transferred to a 3.5 cm dish. A slice anchor was used to hold the tissue in place and immediately overlaid with 5 mL of L-15 media plus 1% FBS in control (DMSO), 70 nM Taxol and 10 μM Nocodazole conditions and imaged immediately.

### Multiphoton microscopy of 3D live cell imaging and analysis of 3D cell migration

Cell migration data was collected by imaging using a custom-built multi-photon laser-scanning microscope (MPLSM) (Prairie Technologies/Bruker) using a Mai Tai Ti:Sapphire laser (Spectra-Physics) to simultaneously generate MPE and SHG to visualize cells and collagen, respectively, at an excitation wavelength of 880 nm with a custom-built temperature-controlled stage insert, as described previously (Ray et al., 2017a, 2017b). Briefly, time-lapse imaging of T cells inside 3D collagen matrix and on tumor slices, was obtained by creating two-channel (T cells and collagen) or three-channel (T cells, collagen, and *KPC*-tissue/*KPCT*-carcinoma cells) Z-stacks of 75 μm depth at 5 μm steps, at each stage position, every 1.5 min over at least 45 min of imaging. Five dimensions (x, y, z, t in two/three channels) time-lapse images were post-processed in Fiji (Schindelin et al., 2012), drift corrected using 3D correction plugin (Parslow et al., 2014) and cell tracking was performed using TrackMate Plugin (Tinevez et al., 2017) in Fiji. Quantification and analysis of tracks was performed as previously described in (Ray et al., 2017a, 2017b). Briefly, cell trajectories were fit to a persistent random walk model (PRWM) (Harms et al., 2005; Othmer et al., 1987) using the method of overlapping intervals (Dickinson and Tranquillo, 1993) using MATLAB (Mathworks) as previously described (Ray et al., 2017b).

Briefly, the mean squared displacement (MSD) for a cell over a time interval *t_i_* was obtained by averaging all the squared displacements *x_ik_* such that

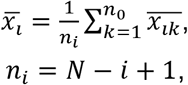

Where *n_i_* is is the number of overlapping intervals of duration *t_i_* and *N* is the total number of intervals. Mathematically, the persistent random walk model can be written as:

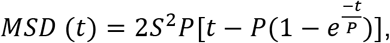

Where *S* is the migration speed and *P* is the persistence time. The motility coefficient is given as:

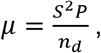

Where *n_d_* is dimensionality. Since the model was fit separately into the three orthogonal directions, we obtained motility, speed, and persistence times for x, y and z directions, therefore *n_d_* = 1 in each case. For total speed of each cell, we took the square root of the squared sum of each speed in the, ◻,◻ and ◻ directions as follows:

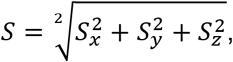

For total motility of each cell, we added the different motilities from each direction as follows:

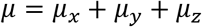

### Statistical Analysis

Multiple groups were compared by ANOVA, followed by the Tukey post hoc analysis. Pairwise comparisons were analyzed using a *t* test. Figure legends indicate which statistical test was performed for the data. Statistical analysis was performed using either KaleidaGraph 4.5.3 (Synergy Software) or Prism 7b (GraphPad Software, Inc). Sample size “N” for each comparison is reported in the corresponding plots (i.e., “N’’ reflects the number of measured individual cells). Number of replicates is 3 or more, unless specified otherwise. Data are shown as box and whiskers diagrams: minimum, first quartile, median, third quartile, and maximum.

## AUTHOR CONTRIBUTIONS

EDT and PPP participated in the initial conceptualization and design of the study. EDT engineered “2.5D” platforms. NJR developed T cell 3D migration assays and live tumor imaging. EDT, NJR, ASZ and PPP designed experiments. EDT, NJR, ACR, MKC, EJP, KY, WSL, ASZ and BRW conducted experiments and data analysis. EDT, NJR, ACR, VP, EAE, EJP, KY, WSR, BRW, BSM, ASZ, and PPP analyzed data. EDT, MKC, EJP, NJR, KY, WSL, and BRW generated unique reagents and their validation. EDT, ASZ, NRJ, and PPP performed quantitative analysis and algorithm implementation. PPP and BSM secured funding. EDT, NJR, and PPP wrote the manuscript. All authors read and contributed comments to the final manuscript. PPP oversaw all aspects of the study.

## ACKNOWLEDGEMENTS

PPP and this work was supported by the NIH (R01CA181385 to PPP, U54CA210190 University of Minnesota Physical Sciences in Oncology Center to PPP, and NIAID training grant T32AI997313 to EJP) and by a Research Scholar Grant, RSG-14-171-01-CSM from the American Cancer Society. This work was also supported by the Randy Shaver Research and Community Fund (PPP) and grants from the UMN Institute for Engineering in Medicine (PPP). ACR was supported by the NIH Intramural Research Program of the National Institute of Biomedical Imaging and Bioengineering. ASZ was supported by the NIH Intramural Research Program of the National Heart, Lung, and Blood Institute. We thank the University of Minnesota Imaging Center (UIC), and UIC staff, particularly Dr. Guillermo Marqués for helpful assistance. The content of this work is solely the responsibility of the authors and does not necessarily represent the official views of the NIH or other funding agencies.

---

**Supplemental Figure 1.**
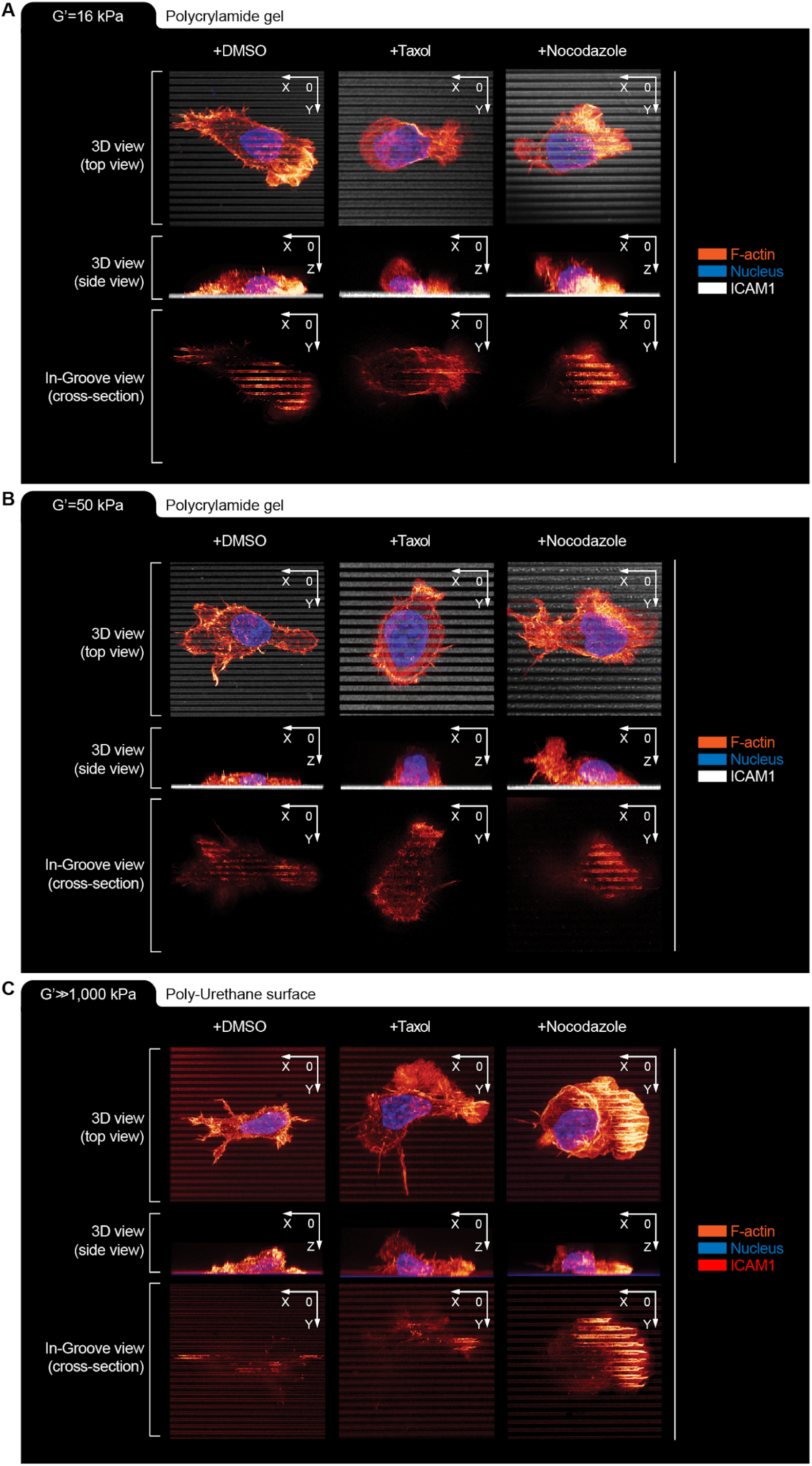
Super-resolution STED 3D microscopy reconstructions of the invasiveness of human cells on nanotopographies of distinct stiffness (related to Figure 1). **(A-C, each panel)** hCD4+ T cell “in-groove” invasiveness: *Left -* control conditions (+DMSO), *Middle* - under microtubules-stabilizing Taxol treatment (+Taxol), *Right* - under microtubules-destabilization conditions (+Nocodazole). **(A)** G’=16 kPa, **(B)** G’=50 kPa and **(C)** G’≫1,000 kPa. **(A-C, each panel)** *Top* - 3D reconstruction view (Top view); *Middle* - 3D X0Z plane (Side view); *Bottom* - “in-groove” X0Y plane view (Cross-section). Note the decrease in T cell “in-groove” invasiveness in control conditions (+DMSO) as the rigidity G’ increases, while Taxol treatment universally decreases “in-groove” invasiveness and Nocodazole treatment universally increases T cell “in-groove” invasiveness. Scale: nanogroove/nanoridge: 800/800 nm.

**Supplemental Figure 2.**
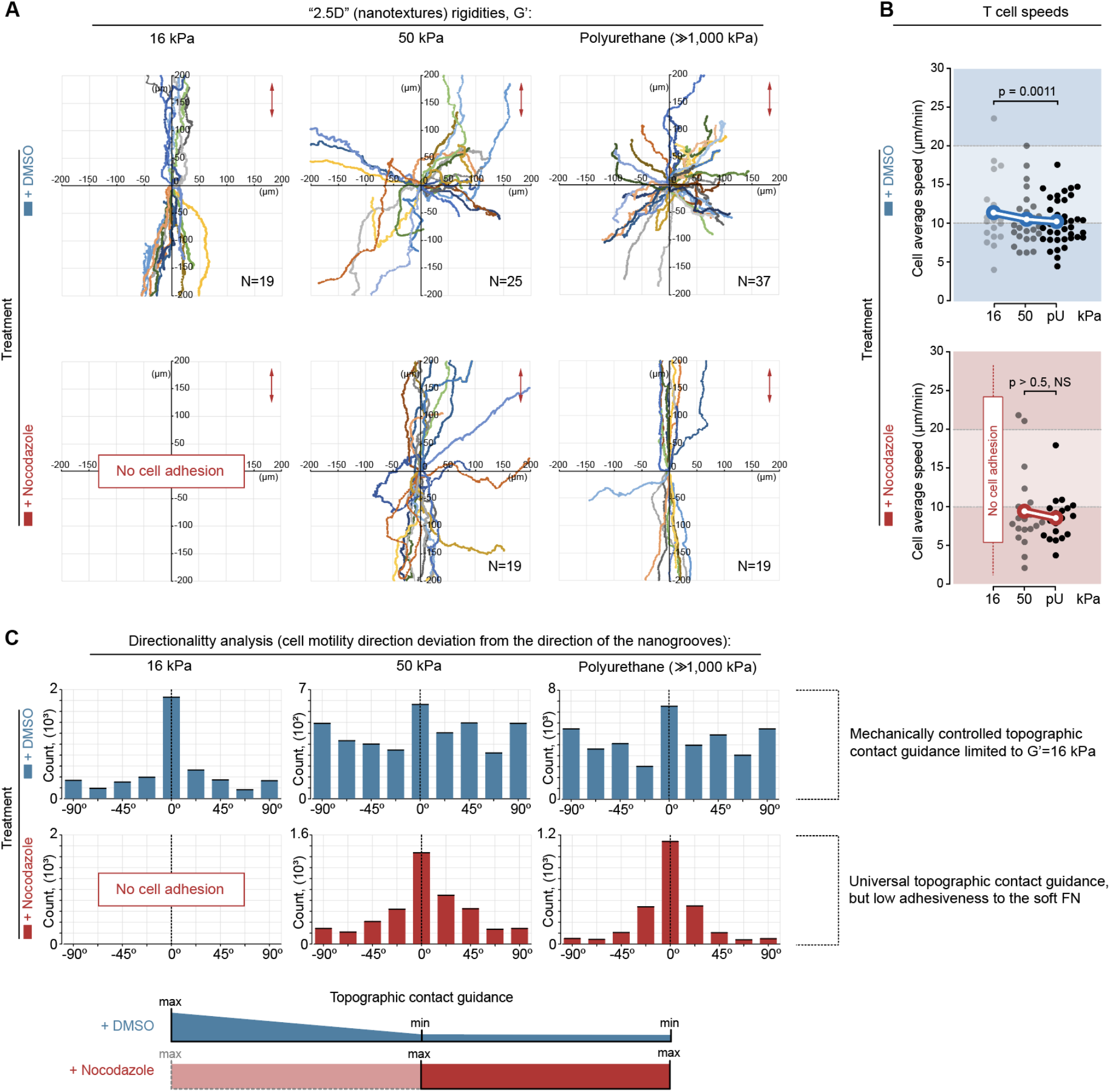
T cell migration on fibronectin nanotextures is stiffness- and microtubule-dependent. **(A)** T cells migration tracks on compliant (G’=16 kPa), intermediate (G’=50 kPa), or rigid (G’≫1,000 kPa) “2.5D” fibronectin (FN) nanotextures, where compliant nanotopographies enhance contact guidance. (*Top to bottom*) T cells migration under control (+DMSO) or Nocodazole conditions, where disassembly of MTs results in the enhanced directed migration across higher rigidities, but limits adhesion on softer architectures. **(B)** Average speeds for T cells migrating on FN nanotextures of various rigidities (16, 50 and ≫1,000 kPa) for control (+DMSO, *top*) and Nocodazole treated conditions. **(C)** Quantification of contact guidance directionality for human T cells migration on FN nanotextures as a function of substrate mechanical rigidity and microtubule stability. Measurements represent frequency distributions of cell-to-nanogroove angles every 10 s step.

**Supplemental Figure 3.**
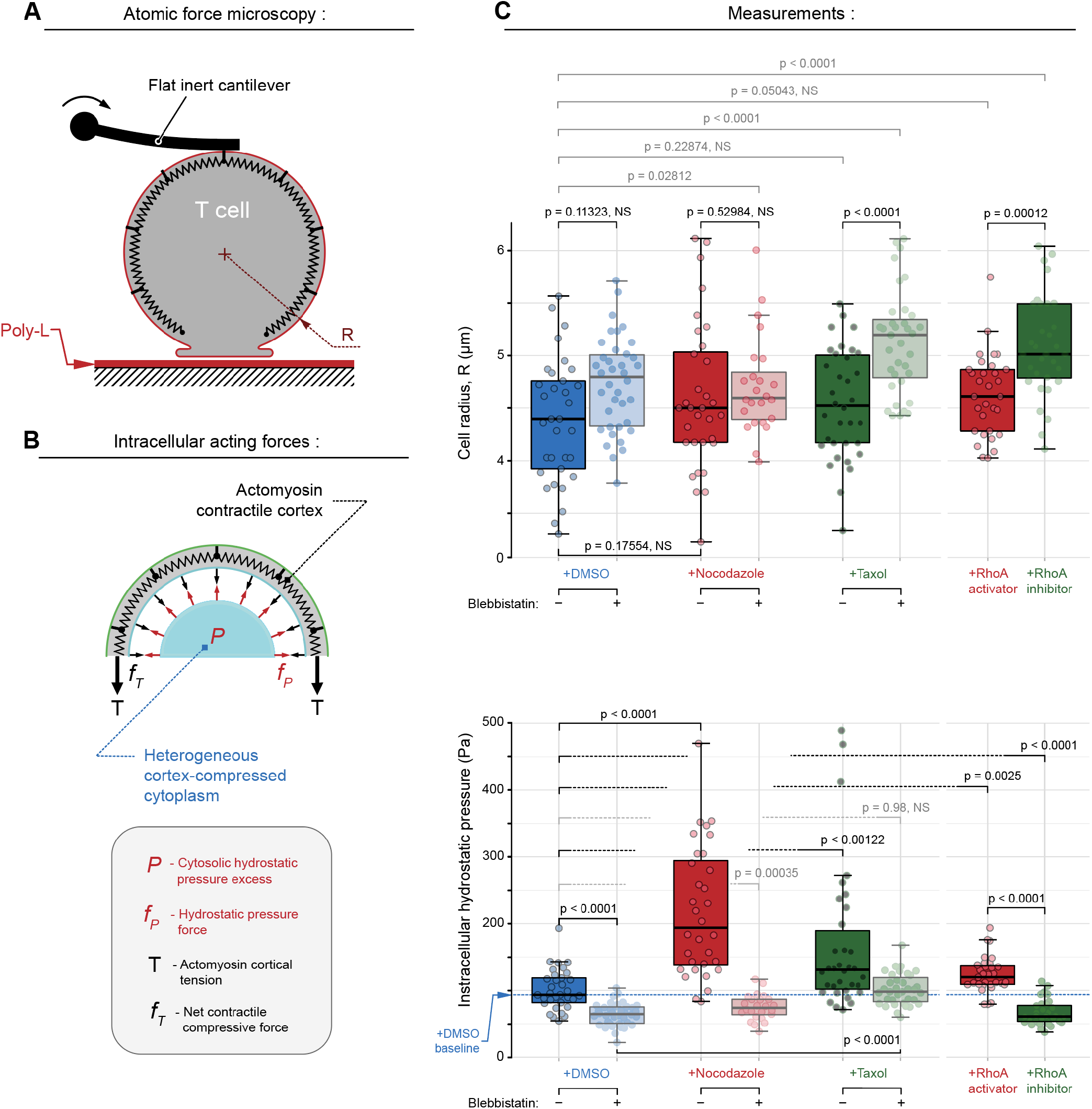
Intracellular hydrostatic pressure in human T cells. **(A)** Schematic diagram of the atomic force microscopy configuration, and **(B)** Schematic diagram illustrating the intracellular force balance in cells under test conditions. The net contractile compressive forces (*f_T_*) generated from the actomyosin cortical tension (*T*) is balanced by the cytosolic hydrostatic pressure excess (*P*). Note that the cortex compressive forces and the corresponding hydrostatic pressure forces (*f_P_*) acts in all directions opposite to each other. Additionally, note that the actomyosin cortex is represented as wiggle lines and the incompressible cytosolic fluid in blue. **(C)** Individual measurements and calculation of the T cell radii and corresponding hydrostatic pressures: *Top* - Human CD4+ T cell radii distribution with and without pharmacological treatments. *Bottom* - Intracellular hydrostatic “cytoplasmic” pressure derived from T cell surface tension and cell shape. The control group and each of the two MT-targeting treatments (i.e., +DMSO, +Nocodazole and +Taxol, *solid colors*) are paired with blebbistatin co-treatment (*pale semi-transparent colors*) to verify the key role of actomyosin contractility in change of intracellular hydrostatic pressure during MT-targeting. Alternatively, direct RhoA activation or inhibition is compared to MT-perturbation results. Both MT disassembly (+Nocodazole) and direct RhoA activation (+RhoA activator) induce significantly increased cytoplasmic pressure in human T cells via increased actomyosin tension, as demonstrated by AFM findings after blebbistatin co-treatment. MT destabilization or direct RhoA activation induced the rise of hydrostatic pressure. Note that taxol-induced stabilization of MTs increases passive (i.e. actomyosin-independent) resistance to compression via a direct mechanical contribution of stabilized MT scaffolds (+Taxol vs. +Taxol+Blebbistatin treatments).

**Supplemental Figure 4:**
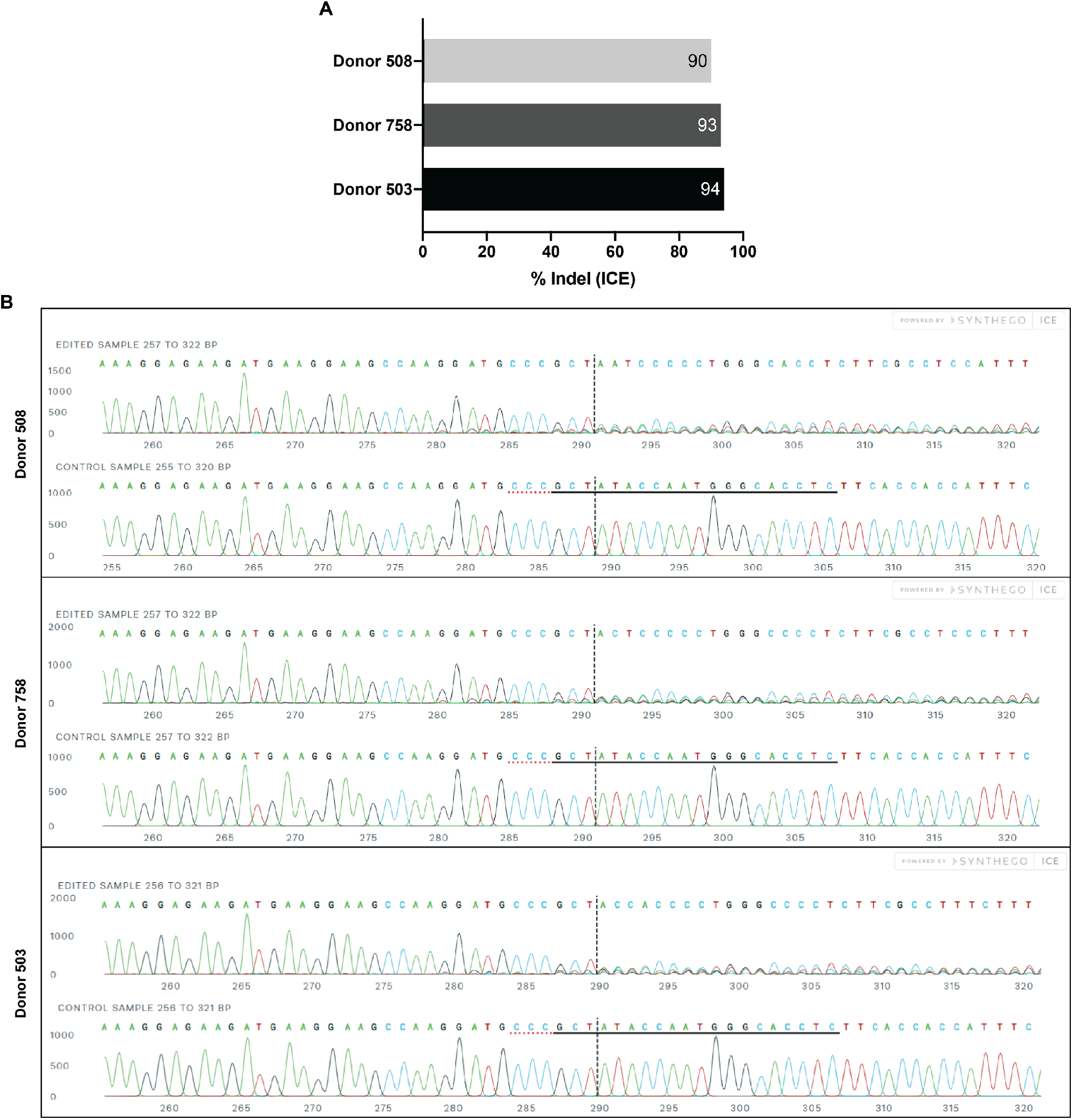
Genotyping of hGEF-H1 KO in human CD4+ (hCD4+) T cells. (**A**) Indel percentage of hGEF-H1 KO in primary T cell lines from 3 separate donors. (**B**) Sanger sequence view showing edited and wild-type (control) sequences in the region around the guide sequence. The hARHGEF2 gRNA sequence is 5’-GAGGTGCCCATTGGTATAGC-3’ while the horizontal black underlined region represents the reverse complementary guide sequence. The horizontal red underline is the PAM site while the PAM CRISPR/Cas-9 recognition sequence is GGG. The vertical black dotted line represents the actual cut site.

